# Integrative machine learning predicts activating kinase mutations for precision oncology

**DOI:** 10.1101/2025.10.14.682355

**Authors:** Yiming Wang, Fangping Wan, Zhangtao Chen, Jonathan Nukpezah, Tom Pan, Kathleen J. Stebe, Cesar de la Fuente-Nunez, Ravi Radhakrishnan

**Affiliations:** Department of Chemical and Biomolecular Engineering, University of Pennsylvania, Philadelphia, PA 19104, USA; Machine Biology Group, Departments of Psychiatry and Microbiology, Institute for Biomedical Informatics, Institute for Translational Medicine and Therapeutics, Perelman School of Medicine, University of Pennsylvania, Philadelphia, Pennsylvania, United States of America; Department of Bioengineering, School of Engineering and Applied Science, University of Pennsylvania, Philadelphia, Pennsylvania, United States of America

**Author notes:** Corresponding authors: Ravi Radhakrishnan,; Cesar de la Fuente-Nunez. YW and FW contributed equally to this work.

## Abstract

Kinases are enzymes that catalyze phosphorylation and play crucial roles in a myriad of cellular regulatory processes and hemostasis. Patient-specific genetic mutations that aberrantly activate kinases can profoundly influence cancer progression and alter drug efficacy. Predicting the impact of such missense mutations across the human kinome on protein function and cellular signaling is therefore a critical step toward personalized targeted therapy. Here, we present Kinome-AI, an integrative machine learning framework that classifies kinase missense mutations as activating or non-activating. Kinome-AI is trained on a rich multi-modal feature set, including residue-level biochemical changes, sequence embeddings from a protein language model, and structural descriptors of kinase–ATP–substrate complexes derived from molecular modeling. Notably, detailed structural features were available for only 21% of mutants; we leverage these as privileged information during training to impute missing structural data for the remaining ∼79. This strategy boosts performance without requiring structural inputs for new (unseen) mutations. The resulting classifier achieves an area under the receiver operating characteristic curve (AUROC) of 0.85 and a balanced accuracy (BACC) of 0.76 across 1,003 mutations spanning 110 different kinases —substantially outperforming existing bioinformatics and general-purpose variant effect predictors. This work provides a robust approach to quantify sequence–structure– function relationships of cancer-driving kinase mutations, paving the way for improved personalized cancer treatment.

**Significance Statement:** In cancer patients, numerous mutations in diverse protein kinases lead to marked differences in disease progression and drug response. Identifying which kinase mutations are activating in individual patients is therefore critical for precision oncology. Drawing inspiration from teacher– student (privileged information) learning, we developed a deep learning framework that integrates structural features from molecular simulations with sequence embeddings from protein language models. This approach enables accurate binary classification of the activation status of kinase mutations. Our study demonstrates how data-driven algorithms can leverage accumulated sequence and structural knowledge of known mutations to predict the effects of novel variants *a priori*. The model, termed Kinome-AI, shows significant promise for incorporation into personalized cancer therapy decision pipelines.

## Introduction

Protein kinases are central regulators of numerous cellular processes, and their dysregulation can lead to cancer initiation and progression [1]. The human kinome comprises 538 kinases, all sharing conserved structural motifs essential for catalyzing the transfer of ATP’s γ-phosphate to serine, threonine, or tyrosine residues of substrate proteins [2-4]. Activating (oncogenic) mutations within the kinase domain can significantly alter catalytic activity, affect regulatory interactions, or influence protein dimerization, localization, and degradation. Such disruptions can imbalance cellular phosphorylation networks and drive oncogenic transformation [5]. For instance, the BRAF V600E mutation constitutively activates the BRAF/MEK/ERK (MAPK) signaling pathway in melanoma [6], and EGFR L858R causes aberrant hyperactivation of EGFR signaling in non-small cell lung cancer [7].

Not all activating mutations reside in the kinase domain; some occur in regulatory regions outside the kinase domain. These allosteric mutations can alter kinase conformation or disrupt autoinhibitory interactions, thereby increasing kinase activity. For example, mutations in the extracellular or transmembrane domains of receptor tyrosine kinases (RTKs) such as HER2 can induce ligand-independent dimerization and constitutive activation in breast cancer [8]. However, here we are focused on those mutations that occur within the kinase domain which harbors a majority of the mutations in human disease, specifically cancers. Given their central role in cancer, kinases have become prime drug targets: hundreds of small-molecule kinase inhibitors are clinically approved to block aberrant signaling [4]. However, the efficacy of these therapies often diminishes due to the vast combinatorial diversity and continual evolution of kinase mutations in tumors [9, 10]. This highlights the need for tools to rapidly characterize whether an individual patient’s kinase mutations will drive oncogenic activity or confer drug resistance.

Computational algorithms have emerged as important aids for predicting the pathogenicity of missense mutations. General-purpose methods leverage sequence conservation and other generic features— for example, FATHMM (Hidden Markov models) [11], Polyphen2 [12], and SIFT [13] use evolutionary conservation, while newer deep learning models like MVP model [14], MutBless [15], MutPred2 [16] and AlphaMissense [17] learn patterns from large variant databases. However, these broad approaches are not specialized for kinases and often show limited accuracy when applied to kinase mutations [10]. Specialized tools targeting kinase variants, such as Kinact [18] and Activark [19], also face significant limitations. These include: (1) the absence of a standardized quantitative approach for determining the activation status of kinase mutants; (2) insufficiency of kinase-only information for indicating kinase activity, since the catalytic process critically involves interactions between Mg^2^LJ, ATP, peptide substrates, and the kinase; (3) limited individual correlation of basic sequence and static structural features (e.g., mutation type, location, protein property predictions, and evolutionary conservation scores) with activation status; (4) training sets biased toward hotspot mutations located within specific kinase domains, such as activation loops and catalytic regions [15]; and (5) small, imbalanced training datasets (N<500), with disproportionate representation of active and inactive mutants [20, 21].

To address these issues, we developed a machine learning framework that leverages an integrative set of residue-level, sequence-based, and structure-based features. We derive these features using state-of-the-art computational tools, including large protein language models, atomistic molecular dynamics (MD) simulations, and molecular docking. In brief, residue-level features capture changes in biochemical properties due to an amino acid substitution (e.g. charge, polarity, size). Sequence-based features include embeddings from a pretrained protein language model, Evolutionary Scale Modeling-2 (ESM2) [22], extracted for both the wild-type and mutant residues in the context of the full protein sequence (as well as the isolated kinase domain sequence). These embeddings encapsulate intrinsic sequence relationships and co-evolution patterns via a self-attention mechanism. Additional sequence features include the identity and chemical class of the wild-type and mutant amino acids, their positions within the protein, and computed changes in physicochemical properties upon mutation.

We encompass structural, energetic, and topological properties of active and inactive conformations for 215 mutants across five kinases, computed from MD trajectories and kinase-ATP-substrate conformations obtained through molecular docking. Inspired by memory-based learning, where previous training examples inform the analysis of new instances, we developed the Kinome-AI model. This model leverages structural information available for 21% of mutants as privileged data to impute missing features for the remaining 79%, enhancing model training without requiring these features during testing. We demonstrate that the classification performance of our framework significantly surpasses other machine learning tools. Further insights were obtained through feature ablation studies and feature importance analyses.

## Results

### Data Curation

Initially, we curated and compiled all available data on 1,003 kinase mutations from literature sources covering over 110 distinct kinases [20, 23]. These kinases were subsequently categorized into eight families based on sequence similarity (Fig. 1A): AGC (including PKA, PKG, PKC families), calmodulin-dependent protein kinase (CAMK), casein kinase 1 (CK1), CMGC (including CDK, MAPK, GSK3, and CLK families), serine/threonine kinase family (STE), tyrosine kinase (TK), tyrosine kinase-like (TKL), and other kinases [24]. Each kinase mutation was assigned a binary label indicating its functional status: activating mutations received a label of 1, while non-activating mutations received a label of 0. These labels were validated through literature-reported data, including in vitro catalytic rates and effects on cell viability (see Methods section).

**Figure 1.**
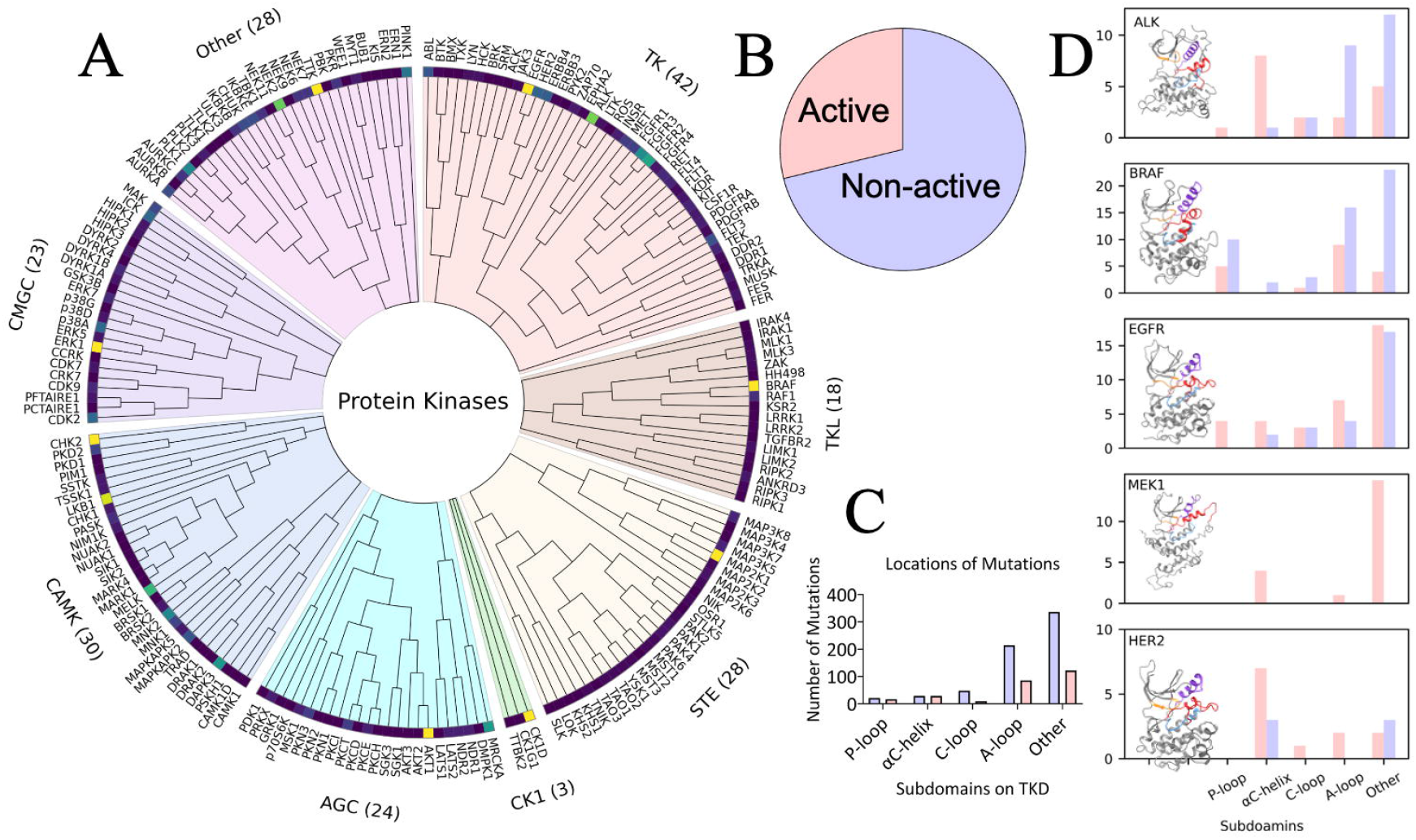
Kinase mutant database. (A) Circular plot displaying 1,003 kinase missense mutations from eight distinct kinase families. (B) Ratio of activating (oncogenic) versus inactivating mutations in the dataset. (C) Distribution of mutation locations across key regions of the kinase domain: the P-loop, αC-helix, C-loop, A-loop, and other regions. (D) Example structures of five kinases in our dataset (one from each of five major families): ALK (PDB: 3LCS [63]), BRAF (PDB: 4MNF [64]), EGFR (PDB: 2GS6 [47]), MAP2K1 (PDB: 3EQD [61]), and ERBB2 (PDB: 3PP0 [65]).

Each kinase domain consists of the following structural regions (or sub-domains): the P-loop, αC-helix, catalytic loop (C-loop), activation loop (A-loop), and other regions (sub-domains). Our dataset contains approximately 30% activating vs 70% non-activating mutations (Fig. 1B). The distribution of mutation locations across these regions is shown in Fig. 1C and 1D. Mutations are not restricted to classical hotspots (such as activation loop positions); in fact, many activating mutations occur outside the canonical catalytic motifs—highlighting that simple rules based on location are insufficient to identify activating variants. Importantly, our dataset selection avoided over-representation of any single hotspot site, resulting in a relatively sparse coverage of the kinase domain sequence space (most mutants are unique single occurrences rather than recurrent hotspots).

### Feature Selection

Regulation of kinase activity is fundamentally linked to the protein’s conformational dynamics [25]. Since a kinase’s three-dimensional conformation is dictated by its amino acid sequence [26] and its molecular environment [27, 28], we reasoned that combining sequence information with structural information would be key to predicting whether a given mutation shifts the kinase toward an active state (thus potentially driving cancer progression).

As summarized in Fig. 2, our feature-generation approach integrates both sequence-based and structure-based features. For sequence-based features, we extract sequence-embedding features from the initial and mutated tokens of kinases using Evolutionary Scale Modeling (ESM-2), a pretrained protein language model. This extraction utilizes both the entire protein sequence and the kinase domain sequence, capturing intrinsic sequence relationships, such as correlations between amino acids, via a self-attention mechanism [22]. Additional sequence-based features include wildtype and mutated residue types, classes, positions, and changes in physicochemical properties resulting from amino acid substitutions, as detailed in a previous report [21]. For further details, refer to the SI Appendix.

**Figure 2.**
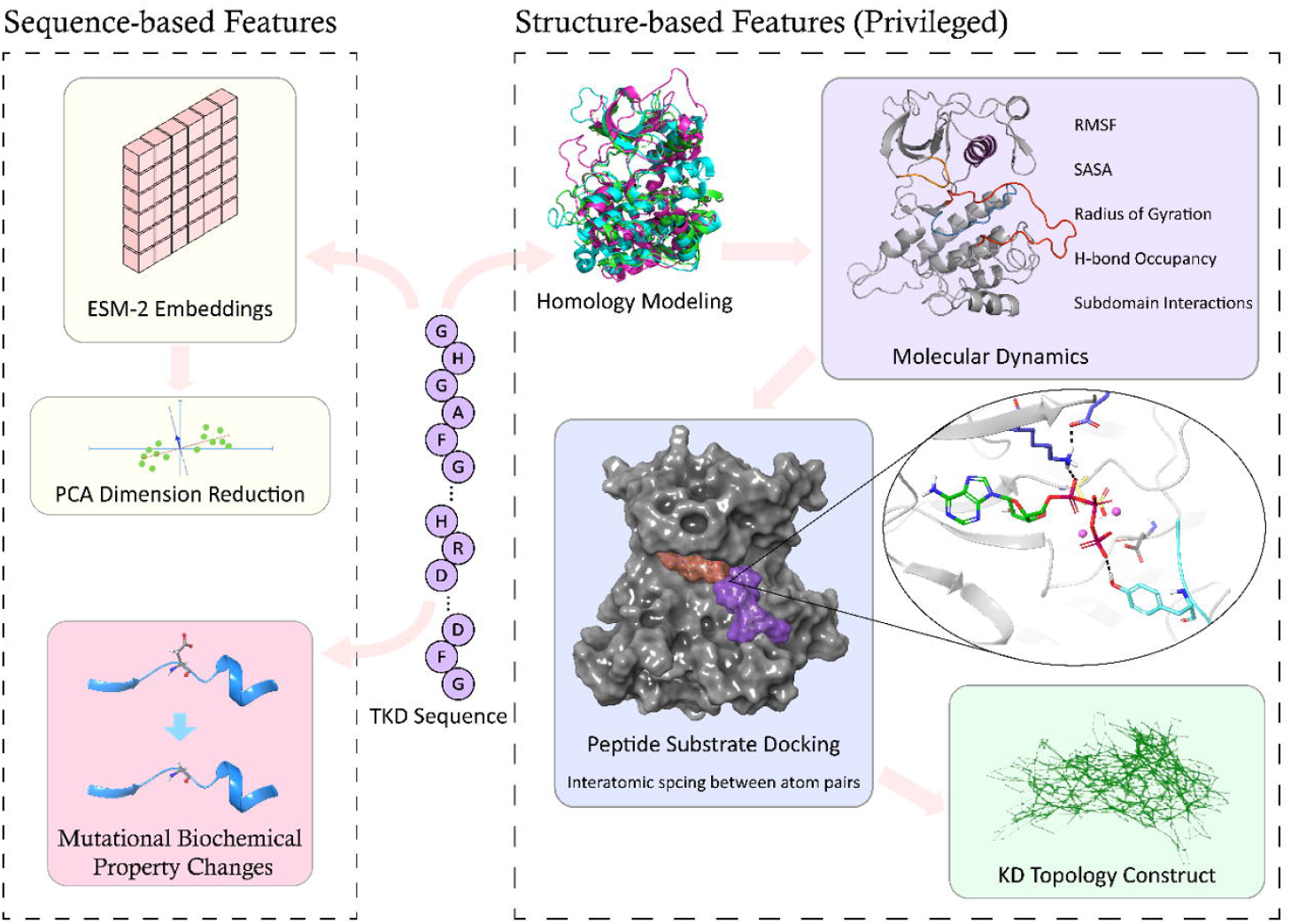
Overview of feature generation for Kinome-AI. (A) Schematic illustration of how sequence-based features (including ESM-2 language model embeddings) and structure-based features (from MD simulations, topology analysis, and docking of ATP and substrate) are derived and integrated for each kinase mutant.

For structure-based sequences, molecular dynamics (MD) simulations have been applied to identify differences in how wildtype and mutated kinases sample inactive and active conformations, such as EGFR [29], ALK [10] and MEK1 [30]. A previous study [10] demonstrated that perturbations in structural properties, like hydrogen-bond occupancy in the αC-helix and activation loop of ALK mutants, effectively indicate activation status. Therefore, besides sequence-embedding features, we computed kinase structural properties from MD simulations of kinase conformations. Specifically, we calculated the mean and standard deviation for root mean square deviation, hydrogen bond occupancy, solvent-accessible surface area, and non-bonded potential energy for domains including the αC-helix, P-loop, catalytic loop, and activation loop from active and inactive conformations across 215 mutants of five kinases (ALK, HER2, EGFR, MEK1, and BRAF). Further details are provided in the SI Appendix and Tables S1 and S2.

To extract geometric information underlying structural variations, we employed a homology tool from algebraic topology to construct a continuous shape known as a simplicial complex from the MD trajectories, reflecting protein geometry across different scales [31, 32]. From these simplicial complex structures, we derived topological features, including the number of connected components, loops, and voids, across various scales. The procedure for topological analysis involves: (i) using molecular docking to identify optimal ATP and peptide substrate binding conformations for each of the five wildtype kinases, (ii) identifying residues within a 6 Å distance from ATP and peptide substrate as catalytic domain residues, and (iii) performing topological analysis to compute connectivity distributions of heavy atoms for residues identified in step (ii).

Finally, structural features for the kinase-ATP-substrate complex were extracted by docking Mg^2+^, ATP, and peptide substrates to each of the 215 kinase mutants in their active conformations obtained from MD simulations. Molecular docking of ATP and peptide substrates with various kinase mutants also offers valuable insights into the kinase-assisted phosphorylation reaction mechanism [33]. From the top-ranked, reaction-competent docking poses, we calculated seven critical pairwise distances involving Mg^2+^, carbons on the HRD-Asp and DFG-Asp sidechains, and the gamma/beta phosphates of ATP, essential prerequisites identified by prior QM/MM studies for phosphorylation reactions [33]. (see Fig. S1 and S2) Additionally, topological features were computed using all non-hydrogen atoms within residues forming the large and small binding pockets identified during ATP-peptide docking, (see Fig. S3).

### Binary Classification

Our goal was to train machine learning models on the above features to classify whether a given kinase mutation is activating (1) or non-activating (0). We distinguish between basic features (the features available for all 1,003 mutants—primarily sequence-based descriptors and kinase family identifiers) and privileged features (the detailed structural features from MD, topology, and docking, which are only available for ∼20% of the mutants). A conventional machine learning approach would require complete data for all features, which would force us to drop or impute the missing privileged features for the majority of the data. Instead, we adopted a *learning with privileged information* approach: during training, the model can utilize the privileged features for the subset of data where they exist, improving its learning; during testing (or when predicting on new mutations), it relies only on the basic features.

We explored several types of classifiers. Traditional models we evaluated included a support vector machine (SVM) and gradient boosting decision trees (GBDTs, including a variant using natural gradient boosting, nGBDT). For these models, we handled missing structural features via k-nearest neighbors (KNN) imputation: for each mutation lacking privileged features, we found its nearest neighbors in the space of basic features and imputed the missing values by averaging the neighbors’ privileged feature values. This yields a complete feature vector that the SVM or GBDT can accept, albeit with some imputation noise.

In parallel, we developed a memory-based neural network (MNN) that inherently manages missing privileged data at the hidden feature level (Fig. 3). This model consists of two subnetworks: a “student” network that processes the basic features (always present), and a “teacher” network that processes the combined basic+privileged features (when available). During training, the teacher network learns an enriched representation of each sample using all features, and the student network learns from those representations even for samples where privileged inputs are absent. Specifically, the MNN uses an attention mechanism to recall patterns from mutants with available structural data and to estimate the latent contribution of the missing features for mutants lacking them. In practice, at each layer of the student network, a set of scaling and shifting factors is predicted (via attention over a memory of privileged-feature examples) to adjust the student’s hidden state, effectively imputing the influence that the privileged features would have had. This approach allows the network to impute missing information internally rather than at the raw input level. The output of the MNN is passed through a differentiable *neural decision forest* classifier: a series of learned binary decision tree nodes that partition the feature space, whose leaf outputs are combined into the final probability prediction. We trained the neural network with a focal loss to handle the class imbalance (down-weighting easy negatives and focusing on harder, rare positives), and we averaged predictions from an ensemble of five MNN models (trained with different random seeds) for stability.

**Figure 3.**
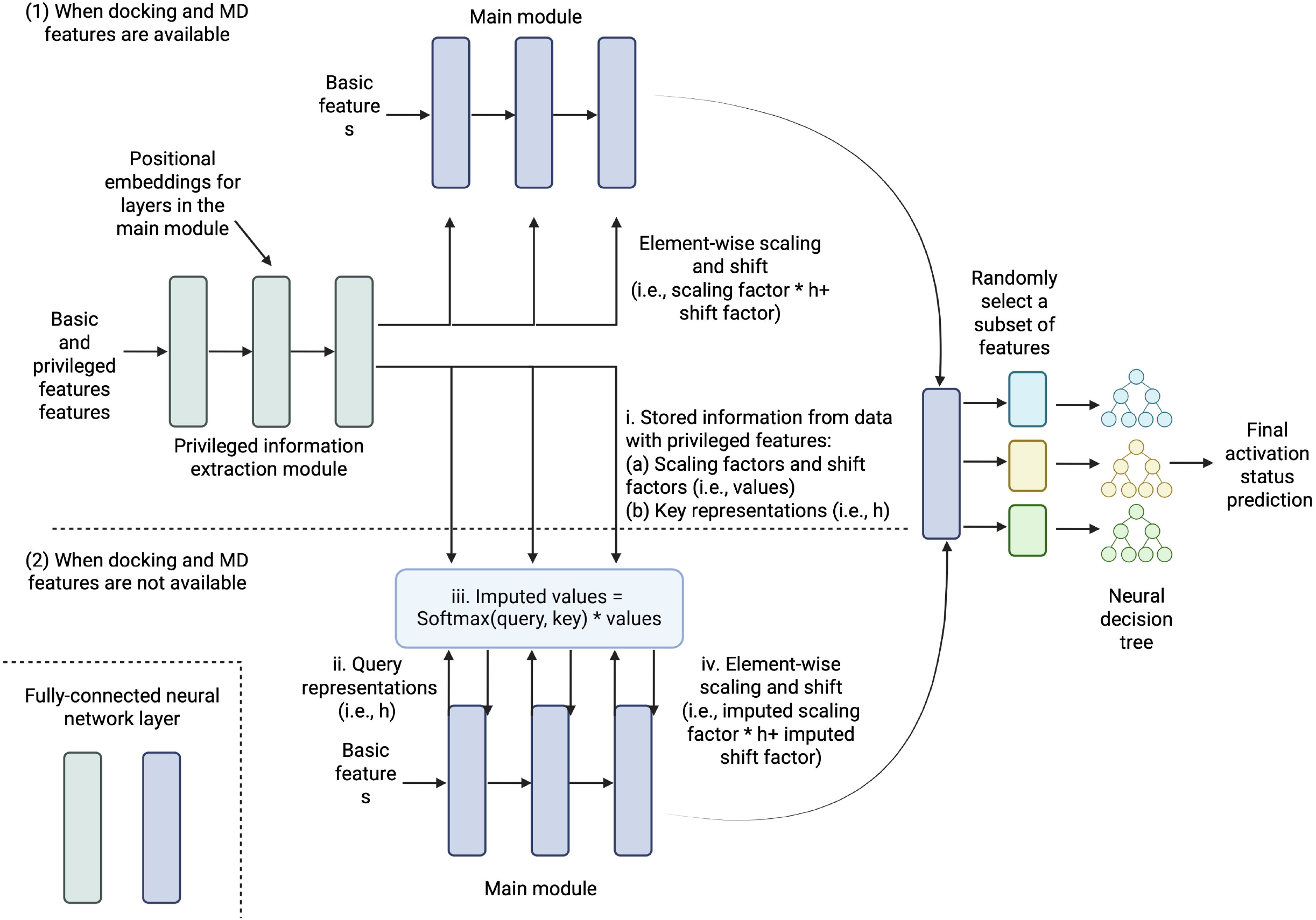
Neural network architecture with privileged learning. Diagram of the Kinome-AI model. A student network processes basic sequence features for all mutants, while a teacher network processes both sequence and structural features for the subset of mutants with available structural data. During training, the model uses an attention-based memory module to adjust the student’s intermediate layers based on teacher representations of similar mutants (privileged information), enabling improved predictions without requiring structural inputs at test time.

### Model Performance

To evaluate the kinase mutation activation prediction performance of our ML models, we conducted five-fold cross-validation repeated five times with varying random seeds. We report performance metrics including the area under the receiver operating characteristic curve (AUROC), balanced accuracy (BACC), and area under the precision-recall curve (AUPRC).

Initially, we trained models without including the ESM-2 sequence embedding features, to assess baseline performance of traditional features. In this setting, the MNN significantly outperformed the others (Fig. 4A,B patterned bars; see also Table S3). The biggest gains from the MNN were observed for mutants lacking structural features (Table S4), indicating that its hidden-layer imputation strategy was effective. For the subset of mutations with full structural data, a GBDT model achieved accuracy on par with the MNN (Table S4), suggesting that when all inputs are present, a well-tuned tree model is quite capable.

**Figure 4.**
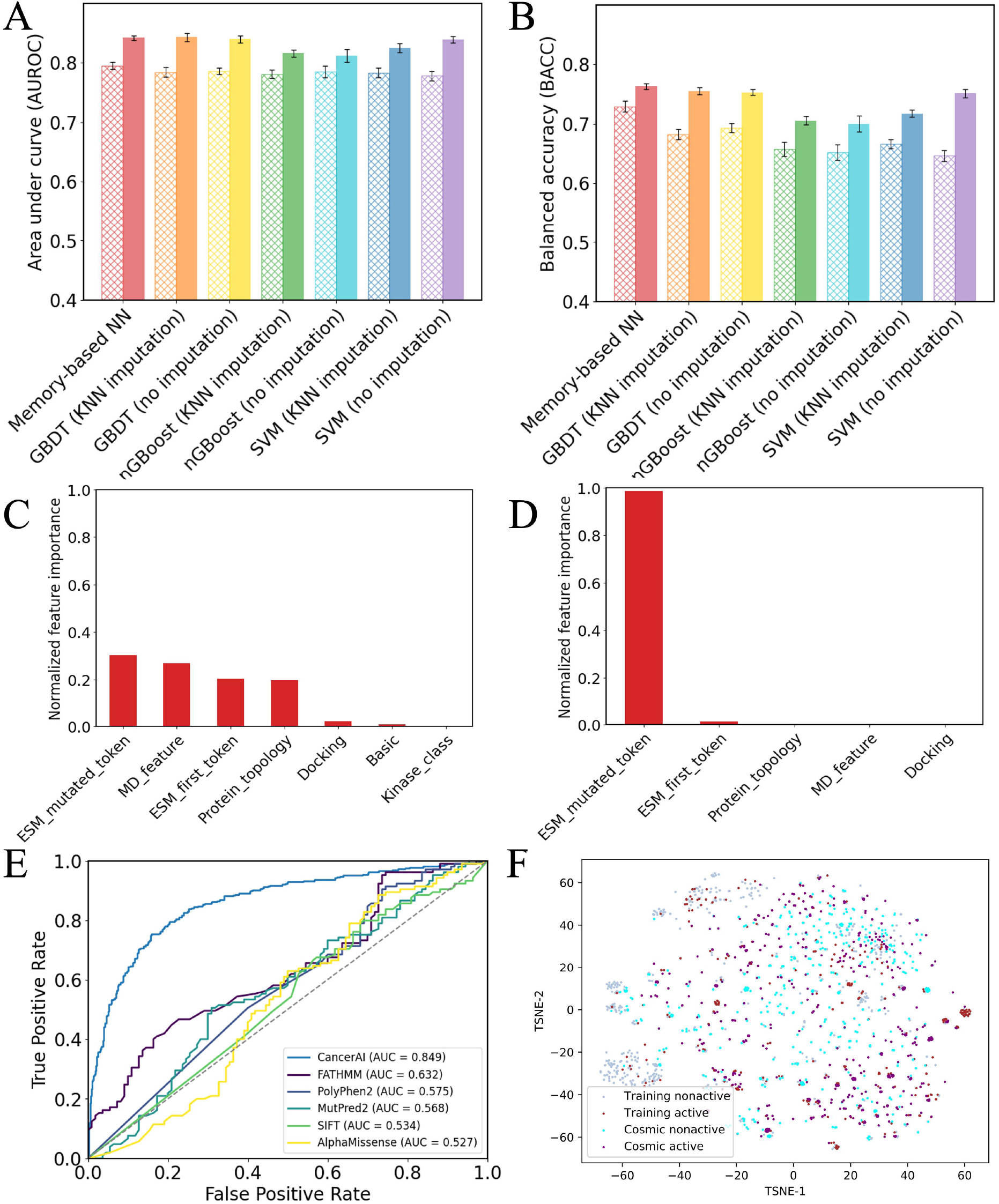
Performance of Kinome-AI and feature importance analysis. (A) Area under the receiver operating characteristic curve (AUROC) and (B) balanced accuracy (BACC) of different models, with (solid bars) or without (striped bars) the ESM-2 sequence embedding features included. The memory-augmented neural network (MNN) outperforms other classifiers, especially when no embeddings are used. (C) Relative feature importance (grouped) in the MNN, computed via integrated gradients attribution. (D) Permutation feature importance (grouped) in the GBDT model. In both models, sequence embeddings (ESM-2) contribute the most to prediction performance, followed by MD-based structural features. (E) ROC curves comparing the Kinome-AI ensemble (MNN+GBDT, black curve) to other variant effect predictors (colored curves) on the task of classifying kinase mutations. Kinome-AI achieves the highest AUROC (∼0.85) (F) t-SNE plot (t-distributed stochastic neighbor embedding [66]) visualizing mutants in embedding space. Points are colored by the model’s predicted class (red = activating, blue = non-activating). Known activating mutants (e.g., ALK *F1174L*, an oncogenic mutation in neuroblastoma cluster in the red region, separate from benign variants.

Incorporating the ESM-2-embedding features into the feature set significantly boosted performance across all models (Figure 4a-b, Tables S5-6). These embeddings provided a powerful representation of sequence context, improving AUROC, BACC, and AUPRC for both the neural and tree-based models. With ESM-2 included, the performance gap between the MNN and classical models narrowed. The MNN still achieved the highest BACC (∼0.76), but tree models were very close and their AUROC/AUPRC were essentially equivalent to the neural network’s (Fig. S4). This suggests that the information captured by large language model embeddings already encompasses much of the signal that might otherwise be gained from explicit structural features. In other words, ESM-2’s embedding of a mutant’s sequence may implicitly reflect some structural and functional context, making the additional MD/docking features somewhat less critical—hence adding those privileged features yielded only modest further improvements once ESM-2 features were present.

Given that two approaches were performing strongly (MNN and GBDT with KNN imputation), we combined them via ensemble learning to maximize reliability. Our final Kinome-AI classifier averages the predicted probability from the MNN and the GBDT, each trained on the full feature set including ESM-2. This ensemble achieved the best overall metrics (AUROC ∼0.85, BACC ∼0.76). Importantly, Kinome-AI substantially outperforms existing tools on this task. Figure 4E compares ROC curves for Kinome-AI vs. several widely used variant effect predictors: FATHMM, PolyPhen-2, SIFT, MutPred2, and the recent deep learning model AlphaMissense. On the kinase mutation dataset, our method shows a markedly higher true-positive rate at all false-positive levels (Kinome-AI’s AUROC was ∼0.85 versus ∼0.65 or lower for the others). Balanced accuracy for AlphaMissense, for example, was only ∼55% on these kinase mutations, consistent with the notion that a general model struggles on this specific subclass of variants. These results underscore the value of a kinase-focused model that incorporates domain-specific biochemical knowledge.

### Feature Importance and Ablation Analysis

We next sought to identify which features or feature classes were most influential in the Kinome-AI model’s predictions. To do this, we conducted feature importance analyses using two techniques: permutation importance for the GBDT model [34] and integrated gradients for the neural network [35]. In both cases, we grouped features into categories for comparison. The feature groups included: basic sequence features (which comprise simple mutation descriptors and kinase family codes), ESM-2 embedding features (for beginner and mutated tokens in the kinase domain), MD-derived structural features, topology features, and docking interaction features.

Figures 4C and 4D show the relative importance of these groups for the neural network and GBDT, respectively. In both models, the ESM-2 sequence embeddings emerge as the single most critical feature class, far surpassing any other group. This aligns with our performance observations that removing ESM features causes a large drop in accuracy. Among the remaining features, the MD-based structural descriptors were the next most important on average, followed by the docking features and basic sequence features. The top 20 individual features (Fig. S5) likewise consisted predominantly of specific dimensions of the ESM-2 embeddings, with a few structural features (particularly some MD features) ranked highly as well.

To validate these findings, we performed ablation studies in which we retrained models after removing certain feature groups (Tables S5, S6). Eliminating the ESM-2 embeddings led to a significant performance drop across the board (e.g., the MNN’s BACC fell from ∼0.76 to ∼0.68), confirming that these embeddings carry essential information. Removing the MD-derived features had a more nuanced effect: for tree-based models and SVM, the change was minimal, suggesting that our KNN imputation of those high-dimensional structural features might not have been very effective (possibly introducing too much noise). However, in the neural network, excluding MD features did degrade performance noticeably, consistent with the idea that the network’s internal imputation (via attention) was making good use of those structural clues. We suspect that the MD features rank high in importance for the GBDT partly because (i) their imputed values were correlated with the target labels (making them appear predictive during training, even if they didn’t generalize as well), and (ii) the limited number of mutants with real MD data might lead the tree model to overweight those features in that subset. The neural network’s approach of imputing in latent space appears to be more reliable, as it could leverage structural signals without as much risk of overfitting to noisy imputed values.

Notably, both the MNN and GBDT performed comparably on our dataset (∼1,000 samples). We anticipate that with larger datasets, the neural approach may have greater headroom for improvement, as it can learn more complex nonlinear interactions and better imputation of missing information, whereas tree-based methods may plateau. As an additional evaluation, we specifically examined ALK kinase mutants as an independent test case. ALK was one of the kinases for which we had a sizable number of mutations and known experimental activities (catalytic rates) from prior work [10]. We trained our model without ALK data, then used it to predict the activation probability of various ALK mutants. Figure 4F (and Fig. S6) shows that the predicted probabilities correlate well with measured kinase activity; mutants known to be highly active (oncogenic) received high scores, while those with low or wild-type-like activity were scored as inactive. Our new Kinome-AI model achieved higher balanced accuracy and a lower false-positive rate on ALK than earlier predictive models we had developed for ALK specifically, Table S7 [10]. This demonstrates that the integrative approach generalizes to specific kinases and can even outperform bespoke models trained for a single kinase.

### Application to Cancer Mutation Data (COSMIC)

A compelling use case for our model is to annotate large cancer genomics datasets to identify likely activating mutations. We applied

Kinome-AI to missense variants recorded in the COSMIC database [36] for several key cancer-associated kinases: ALK, BRAF, HER2 (ERBB2), MAP2K1 (MEK1), and EGFR. These kinases are frequently mutated in tumors and have well-characterized roles in oncogenic signaling. We compiled all unique missense mutations in their kinase domains from COSMIC and used our ensemble model (MNN+GBDT) to predict their activation status probabilities.

The results are summarized in Figure S7. For each kinase, we observed that a large fraction of possible residues in the kinase domain have been found mutated in patient tumors (over half of the positions in some cases), illustrating the breadth of genetic variation cancer cells explore. Our model provides a score for each variant; for visualization, we binned variants into predicted activating vs non-activating. The predictions for different kinases reveal some kinase-specific trends—e.g., certain regions in BRAF and EGFR are enriched for predicted activating mutations—consistent with known hotspots, but also suggesting new potentially activating sites. We note that the MNN and GBDT components of the ensemble sometimes differed in their confidence for specific variants (likely due to their different architectures), but the overall ensemble averaged these to provide a consensus. A systematic comparison of the model performance across all eight kinase families (Table S8) indicates that the ensemble approach is robust across diverse kinase types. These predictive maps of mutational activation can guide experimental prioritization: for instance, novel mutations that our model flags as likely activating could be tested in lab to confirm their impact and potentially inform treatment (especially if found in a patient’s tumor).

## Discussion and Conclusions

Understanding which mutations drive kinase activation is crucial for interpreting cancer genomes and tailoring therapies, yet it remains a challenging problem at the intersection of structural biology and genomics. While recent advances like AlphaFold have revolutionized protein structure prediction [37], determining how a mutation alters protein *function*—particularly for enzymes with complex regulation like kinases—requires going beyond static structure to capture dynamics and molecular interactions. Kinase activation involves intricate conformational changes and assembly of a multi-component complex (kinase, ATP, Mg^2^□, substrate peptide), a process that is highly dynamic and still not fully understood. The challenge is compounded by the sheer number of possible mutations: indeed, more than half of the amino acid positions in many kinase domains have been observed to mutate in cancers (Fig. S7) [36]. Outside a few well-known hotspots (e.g., activation loop positions [38], the functional impact of most kinase mutations remains uncharacterized.

Our work demonstrates that integrative computational modeling can help fill this gap. We show that by combining large-scale sequence information with structural biophysics, we can predict mutation impact with a high degree of accuracy. Kinases have a rugged energy landscape with multiple conformational states [10], and oncogenic mutations often work by stabilizing an active state or destabilizing an autoinhibited state [30]. Traditional structure prediction tools (like AlphaFold [37]) output a single static conformation and do not indicate which state a mutant might favor, limiting their use for functional predictions. Similarly, generic machine learning models trained on broad datasets (e.g., AlphaMissense [17]) might capture general features of deleterious vs neutral mutations, but they often miss subtleties of *specific* protein families; as we saw, AlphaMissense and others underperformed on kinase activation prediction (Fig. 4E). This highlights the need for fine-tuned models that incorporate domain-specific knowledge and data.

By focusing on kinases and leveraging mechanistic insights (such as the importance of ATP-binding pocket geometry, or the dynamic coupling of the αC-helix and activation loop, our model gains both accuracy and interpretability. The dominance of sequence embeddings in our feature importance analysis suggests that the evolutionary history encoded in protein sequences is extremely informative about function; however, the additional boost we obtained from structural features, and the improved generalization on ALK mutants, indicate that there is complementary value in those biophysical descriptors. The privileged learning approach allowed us to use those valuable but sparsely available structural features without sacrificing the breadth of our dataset.

In summary, we developed a robust machine learning framework, Kinome-AI, that integrates sequence-based and structure-based features to classify kinase mutations as activating or non-activating. Our model achieved high overall performance (AUROC 0.85, BACC 0.76 on 1,003 variants from 110 kinases) and outperformed a range of existing variant effect predictors. Beyond performance metrics, Kinome-AI offers biological insights: for example, it highlighted which molecular features (like language model-derived sequence motifs) are most predictive of activation, and it successfully identified which new ALK mutations are likely to be oncogenic— predictions that were used to guide patient-specific therapy in a recent neuroblastoma clinical trial [9]. These results illustrate the potential of combining data-driven AI with mechanistic modeling to advance precision oncology. Our approach is not limited to kinases; it could be generalized to other protein families where some structural or high-resolution data exist alongside larger sequence datasets. For instance, similar strategies could be employed to predict functional effects of viral mutations on infectivity or immune escape, aiding vaccine design [39].

Despite its strengths, our study has several limitations warranting future improvement: (1) The “activation” labels for kinase mutations in our dataset were compiled from diverse experimental sources (in vitro enzyme assays, cell-based signaling readouts, clinical observations of drug response). These different assays may not perfectly agree, introducing noise. In the future, assigning confidence scores to labels or focusing on more uniform criteria could improve model training; (2) Our model considers the kinase domain in isolation, but real cellular kinase activity is influenced by broader context—e.g., the requirement of EGFR to form an active dimer [40], the role of co-factors like BAK for EFR signaling [41], or scaffold interactions like KRAS binding to MEK1 [42]. Incorporating such context (through protein–protein interaction features or cell-specific data) could further refine predictions; (3) We limited our analysis to mutations in the kinase catalytic domain. However, mutations in regulatory regions (N-terminal or C-terminal lobes, docking motifs, accessory domains like FHA domains [43] can also alter kinase activity by modulating regulation, localization, or protein–protein interactions. Expanding the model to include these regions—and their context—would broaden its applicability; (4) Our structural features from MD are dependent on simulation length and quality. Some subtle effects of mutations might require longer sampling or advanced techniques (e.g., enhanced sampling, free energy calculations) to capture reliably. As computational resources improve, more exhaustive simulations of kinase mutants (including full kinase–ATP–substrate complexes in solution) could provide even more informative structural descriptors; (5) Finally, increasing the dataset of kinase mutations with high-confidence functional labels will naturally improve the model. There likely remain many uncharacterized kinase mutations. In parallel, utilizing modern representation learning (for example, training task-specific transformer models on protein structures or dynamics) could automatically generate richer features than the handcrafted ones we used. Such developments could further boost predictive power and generalization. Despite these limitations, it is remarkable that our integrated algorithm—combining sequence, structural, and biophysical features—can already predict the functional activation state of kinase domain mutations with approximately **75% accuracy**. This level of performance, achieved despite the inherent noise and complexity of biological data, underscores the robustness and generalizability of our framework, and highlights its potential as a powerful tool for prioritizing and interpreting functional mutations in large-scale cancer genomics and precision medicine efforts.

In conclusion, our work presents a powerful approach for predicting the functional impact of kinase mutations by marrying data-driven machine learning with mechanistic modeling. As genomic sequencing becomes routine in oncology, tools like Kinome-AI can help interpret mutations in the context of signaling networks and suggest which ones might be driving a patient’s cancer. This can inform precision treatment choices—such as selecting targeted kinase inhibitors when an activating mutation is present, or avoiding certain drugs if a resistance mutation is detected. Moreover, the general strategy of *privileged integrative learning* can be applied to other biomedical challenges where limited high-resolution data exist. We envision a future where such computational models are regularly used alongside experiments to accelerate discoveries and tailor interventions in cancer and beyond.

## Materials and Methods

### Machine Learning Models

We implemented several machine learning approaches to predict kinase activation states, specifically utilizing support vector machines (SVM), gradient boosting decision trees (GBDT), and natural gradient boosting decision trees (nGBDT) as baseline models. Due to the incomplete availability of docking and molecular dynamics (MD) simulation features, termed privileged features, K-nearest neighbor imputation was used to handle missing values. The SVM, GBDT, and K-nearest neighbor algorithms were implemented using scikit-learn.

A novel memory-based neural network model was introduced to enhance prediction performance. This model integrated both basic features (available for all samples) and privileged features (only partially available). The neural network involved several stages. Initially, a fully connected neural network (i.e. a student network) processed the basic features through multiple layers of nonlinear transformations. Subsequently, a separate neural network (i.e., a teacher network) processed a combined vector of basic and privileged features, which was further adjusted using sinusoidal positional encoding. This encoding generated scaling and shifting vectors dependent on the neural network layer index and adjust the vector features generated from the teacher network via Feature-wise Linear Modulation scheme (FiLM) [44]. These vectors were then used to adjust the original hidden features in every layer of the student network, effectively integrating privileged information.

To enable predictions for samples lacking privileged features, a memory-based approach was adopted. Intermediate features derived from privileged information were stored and then employed in an attention mechanism [22] to impute scaling and shifting factors for new samples. Each neural network layer in our context comprised linear transformations, layer normalization, dropout, and Gaussian Error Linear Unit (GELU) activation functions [45]. Shared weights were employed across layers, excluding the final layer, to enhance consistency.

The prediction output of the neural network was integrated with a Deep Neural Decision Forest approach. Here, a binary decision tree framework was employed, with internal nodes routing samples based on randomly selected features transformed by linear layers and sigmoid activation. Leaf nodes represented binary classification decisions, and final predictions were computed as weighted averages of leaf node logits, normalized by softmax probabilities. Training utilized focal loss to mitigate class imbalance issues, and robustness was ensured by averaging predictions across five model instances trained with different random seeds.

Performance evaluation and hyperparameter tuning were systematically conducted through repeated five-fold cross-validation (CV). Model performance was measured using balanced accuracy (BACC), area under the precision-recall curve (AUPR), and area under the receiver operating characteristic curve (AUROC). Hyperparameters for all models underwent extensive grid searches to optimize AUROC performance. Parameters such as tree depth, number of trees, regularization strengths for SVM, and neural network hyperparameters (dropout rates, decision tree depth, number of trees, and training epochs) were comprehensively evaluated.

Feature importance was assessed through permutation importance for GBDT models, while integrated gradient analysis was applied to neural network models. This involved comparing mutant kinase features against their wild-type counterparts as baselines, providing insights into the relevance of individual features in predicting kinase activation. For details, see SI Appendix.

### Molecular Dynamics (MD) Simulations

TKD variants with point mutations and their corresponding active conformation homology models were constructed using MODELLER [46]. For the wild-type EGFR structure, missing residues in the crystal structure (PDB: 2GS6 [47]) were completed using PyMOL, followed by energy minimization. Molecular dynamics (MD) simulations of 215 kinase mutant proteins were conducted using GROMACS 2020.6 [48] with TIP3P water [49] as the solvent in a periodic box environment and utilizing the CHARMM27 force field [50]. Sodium and chloride ions were added to neutralize the system at an ionic strength of 0.1 M. The simulation box dimensions were approximately 9×9×9 nm^3^, and periodic boundary conditions were applied in all three spatial directions. A cutoff distance of 1.0 nm was employed for both van der Waals and Coulombic interactions, with the Particle Mesh Ewald (PME) method used for long-range electrostatic calculations [51]. Initially, energy minimization was performed using the steepest descent method, followed by equilibration through isothermal-isobaric (NPT) simulations for several nanoseconds at 300 K and 1 bar, using the Berendsen thermostat [52] and Parrinello-Rahman barostat [53], respectively. During the production stage, the temperature was maintained using the stochastic velocity rescaling thermostat [54]. A timestep of 2 fs was used, with trajectories recorded every 100 ps. Bonds involving hydrogen atoms were constrained using the LINCS algorithm [55]. Each system underwent a total simulation duration of 100 ns. For details, see SI Appendix.

### Molecular Docking

ATP^4-^ and peptide substrates were sequentially docked onto the relaxed active kinase domain (KD) structures of ALK, BRAF, EGFR, HER2, and MEK1, extracted from the final frame of molecular dynamics (MD) simulations. These structures contained two Mg^2^□ ions and were docked using Schrödinger’s (version 2020.3) Induced-Fit Docking (IFD) [56] and Glide protocols [57]. Positions of the Mg^2^□ ions were modeled based on Mn^2^□ ions from a ternary PKACA-ATP-Peptide complex (PDB ID: 1ATP [58]), following alignment of receptor KDs to this reference structure. Additionally, water molecules within 10 □ of the Mg^2^□ ions were retained throughout the docking simulations [58]. Proposed peptide substrates for receptor tyrosine kinases (RTKs) were derived from their respective autophosphorylation sites, specifically Y1278 within the Y’XXX’YY motif on the activation loop (A-loop) of ALK TKD and Y1092 (PEYINQ), a major autophosphorylation site located in the cytoplasmic tail region of EGFR [33, 59]. As HER2 heterodimerizes with EGFR for activation, the EGFR TKD peptide substrate was also utilized for HER2 [60]. For the serine/threonine protein kinases BRAF and MEK1, peptide substrates were selected from the phosphorylation sites in the A-loop regions of their downstream effectors. Specifically, peptide substrates were modeled based on S218 (IDSMA, PDB ID: 3EQD) of MEK1 and T183/Y185 (FLTEYVA, PDB ID: 5UMO) of ERK2 [61, 62].

For details, see SI Appendix.

## Data Availability

All data supporting the findings of this study, including the curated kinase mutation dataset and engineered feature values, are available in the Github repository: https://github.com/wyiming0318/CancerAI. The repository also contains code for training and evaluating the models, as well as scripts for performing the molecular simulations and feature extractions described above. Additional data are provided in the Supplementary Information.

## Acknowledgments

We thank members of the Radhakrishnan and de la Fuente research groups for insightful discussions. We gratefully acknowledge partial funding support from DOE grant 583468 AMND 1/DE-SC0022240, NIH grants R35GM138201, CA250044, and CA244660, and Defense Threat Reduction Agency (DTRA) grants HDTRA11810041, HDTRA12110014, and HDTRA12310001. Computational resources were partially provided by ACCESS under grant MCB200101. All authors declare no conflicts of financial interest with respect to this study.

