## Supplementary material for "Integrative machine learning predicts activating kinase mutations for precision oncology": SI methods, figs, tables

Kathleen J. Stebe^a^, Cesar de la Fuente-Nunez^acd*^, Ravi Radhakrishnan^abd*^

a Department of Chemical and Biomolecular Engineering, University of Pennsylvania, Philadelphia, PA 19104, USA

b Penn Institute for Computational Science, University of Pennsylvania, Philadelphia, PA 19104, USA

c Machine Biology Group, Department of Psychiatry and Microbiology, Institute for Biomedical Informatics, Institute for Translational Medicine and Therapeutics, Perelman School of Medicine University of Pennsylvania, Philadelphia, PA 19104, USA

d Department of Bioengineering, University of Pennsylvania, Philadelphia, PA 19104, USA

+ YW and FW contributed equally to this work

Cesar de la Fuente-Nunez,

**This file includes:**

Supporting information text

Figures S1 to S8

Tables S1 to S8

SI References

**Supporting Information Text**

**Kinase mutation dataset and their activation status assignment.** Kinase mutations used in this study includes those used in the previous work [Patil PNAS] (link: <https://github.com/kksuresh25/Cancer-AI>) with a few labels corrected and some additional kinase domain mutations taken from the Precision Oncology Knowledge Base (OncoKB) (<https://www.oncokb.org>) curated by Memorial Sloan Kettering Cancer Center New York. [1, 2] The activation status label of kinase mutations were annotated based on in vitro biochemical or in vivo cell assays reported in the literature. These experimental methods mainly consist of enzyme kinetics studies, immunoblotting, cell viability/proliferation, colony formation and focus formation assays that characterize the kinase catalytic rate (k_cat_), fold change in kinase activity or substrate phosphorylation, autophosphorylation, and downstream effector activation of the mutant. To ensure the robustness of activation status labeling, all selected mutants were expressed in the cell assays individually and mutant variants with conflicting in-vitro characterizations were not included. In addition, reported results of the in-vitro experiments were examined to ensure that there is quantitative difference (normalized fold change > 2) between the wide type and the mutant for activating mutations.

**Generation of basic features**. Here the 59 basic bioinformatics features includes the binarized assignment (1/0) of location of the mutations in terms of 4 subdomains (e.g. nucleotide binding loop, αC-helix, catalytic loop, and activation loop), the 20 types and the 5 classes (e.g. aliphatic, acidic, basic, aromatic and polar) of both wild-type and mutated residues, and the five biochemical property changes upon mutation, including the Eisenberg hydrophobicity, the free energy of solvation, van der Waals radius, charge, and polarity. All the five biochemical features were normalized by the range of the properties, according to the formula ${\Delta P}_{MUT}=\frac{P_{WT}-P_{MUT}}{Max(P)-Min(P)}$.

**Generation of sequence-embedding Feature.** The evolutionary scale model (ESM-2) model is a large protein language model trained to predict the identity of amino acids that have been randomly masked out of protein sequences. Here we extract the first token and mutated token of sequence-embedding features from the ESM-2 model using both the whole protein sequence and kinase domain sequence, which encodes general structural information via a self-attention mechanism. Amino acid sequences of mutated kinases were created with a string manipulation Python script that substituted the wild type amino acid with the mutant amino acid at the location of the point mutation, based on our curated mutant database. The kinase domain sequence was also extracted from the whole protein amino acid sequence for downstream use. ESM-2 embeddings were generated using the 33 layer, 650 million parameter ESM-2 variant, which accepts an amino acid sequence as input and produces embeddings of size 1280 per amino acid [3]. The model output also includes embeddings corresponding to a beginning and end of sequence token. Two sets of embeddings were generated from wild-type kinase domain sequences and mutant kinase domain sequences, respectively. The ESM-2 model was downloaded and utilized according to the instructions from [https://github.com/facebookresearch/esm](https://urldefense.com/v3/__https:/github.com/facebookresearch/esm__;!!IBzWLUs!VPug2am5KCBW5ArdXKObPzQc6ap7GxHV2VpFoy-_HvfXsjFHgVbMrXOve6sHedukqInI5kud7zdFLaaFd0r_72Sh$). All embeddings were generated with a Nvidia GeForce RTX 3070 GPU.

Dimensionality reduction was applied to the first token (beginning of sequence token) using a scikit-learn pipeline of standard scaler (applied feature-wise across the samples) followed by principal component analysis (PCA) that reduced embedding dimension from 1280 to 100. To reduce dimensionality of first token embeddings, pipelines of standard scaler followed by PCA were fitted to the first token embeddings of the wild type kinase domain individually. These fitted pipelines were then used to reduce dimension of mutant whole protein and mutant kinase domain first token embeddings respectively. A similar dimensionality reduction technique was applied to the mutation site embeddings. However, these standard scaler and PCA pipelines were fitted separately to all the amino acid embeddings (excluding beginning of sequence and end of sequence tokens) created from wild type kinase domain sequences. These pipelines were used to reduce dimension of the embedding at the mutation site for the kinase domain. They were also used to reduce dimension of the respective wild type kinase domain embedding corresponding to the mutation location, so the wild type feature can be compared to the mutant feature in the neural network downstream. We term these reduced dimension embeddings as the protein language model (PLM) features.

**Molecular dynamics (MD)** TKD variants with point mutations and the homology models of the corresponding active conformations were constructed using MODELLER [4]. For wild-type EGFR structure, missing residues in the crystal structures (PDB: 2GS6) were filled using PyMoL. [5] Followed by a minimization of the structure, MD simulations of 221 TKD models were performed using GROMACS 2020.6 [6] with TIP3P [7] water as solvent implemented in a periodic box with the CHARMM27 force field [8]. Sodium chloride were used to neutralize the system with salt concentration of 0.1M. Simulation box size is around 9x9x9 nm^3^, with periodic boundary condition applied in all three directions. Cutoff distances for both van der Waal and columbic potential are 1.0 nm. Particle mesh Ewald (PME) summation method is applied for long-range electrostatic interaction. [9] First, steepest descent method was applied for energy minimization of initial configuration, which was then equilibrated by isothermal-isobaric (NPT) simulation for a few nanoseconds at temperature of 300K and pressure of 1 bar using the Berendsen thermostat [10], and the Parrinello-Rahman barostat [11], respectively. At production stage, the temperature is maintained using the stochastic velocity rescaling thermostat. [12] A time step of 2 fs is used and simulation trajectory is saved every 100 ps. Bonds with hydrogen atoms were constrained using the LINCS algorithm. [13] The total simulation time for each system is about 100 ns.

**Generation of MD features** Time-averaged conformational MD features were calculated from all frames in the last 50 ns of the trajectories for the active and inactive conformations using GROMACS 2020.6. [14] The MD features recorded the differences between the mutants and the wild-type in terms of root-mean-square-fluctuation (RMSF), solvent-accessible surface area (SASA), and radius of gyration. Pairwise interactions between different subdomains and the interactions within each of the four subdomains (including αC-helix, P-loop, catalytic loop, and activation loop domains) were calculated. Differences between the corresponding residues on the wild-type and mutant were taken before being accumulated for each mutant, producing a total of 74 features. The RMSF, SASA, and radius of gyration of each subdomain in the system, and the pairwise interaction energies between subdomains and within each subdomain were calculated using GROMACS.

**Hydrogen bonding occupancy calculation.** Hydrogen bonds were determined using python package MDAnalysis 1.0.1. [15] For each trajectory, 500 frames sampled from the last 50 ns of the MD simulation were used for the calculations. Step 1. For hydrogen bonding (HB) occupancy of wildtype and mutant ALK protein, first calculate average number of HBs formed in each residue ($O_{WT, i}$ and $O_{MUT, i}$) from both the αC-helix domain (residues 1158-1176) and activation loop (residues 1268-1291) from the last 50ns of two independent MD simulation trajectories. Note: the HB is considered to be formed if 1). distance between hydrogen acceptor and hydrogen donor is less than 0.32nm and 2). the acceptor-hydrogen-donor angle is larger than 150°. In addition, HB is excluded if the salt bridge is formed between the H-donor and H-acceptor. Step 2. Calculate the HB occupancy difference between the mutant and wildtype ALK protein as $\Delta_{MUT, i}=O_{MUT, i}-O_{WT, i}$ for each residue within the αC-helix and activation loop domains. If the condition $\left| \Delta_{MUT, i} \right|>0.75$ is satisfied, then $\Delta_{MUT, i}$ will be added to the accumulated HB occupancy, $A_{MUT}=\sum\Delta_{MUT, i}$.

**Generation of topological features.** Our approach analyzes protein structures by converting them into mathematical frameworks called simplicial complexes. [16] These complexes represent the protein's 3D structure through a series of connected geometric components: points (amino acid positions), edges (connections between points), triangles, tetrahedra, and higher-dimensional analogs. We create these simplicial complexes progressively by connecting points that fall within increasing distance thresholds. This creates a "filtration"—essentially a series of nested simplicial complexes that capture the protein structure at multiple scales of resolution.

Persistent homology tracks topological features across this filtration. Specifically, it identifies:

1. Connected components (0-dimensional features)

2. Loops or tunnels (1-dimensional features)

3. Cavities or voids (2-dimensional features)

As we analyze the protein structure across different distance thresholds, these topological features appear and disappear. The "lifetime" of each feature—when it forms and when it collapses - is recorded in persistence diagrams. These diagrams serve as topological signatures, with each bar representing a specific topological feature and its duration across the filtration. From these persistence diagrams, we extract the specific amino acid residues (Cα atoms) that form these topological features. [17] We construct a network where Cα atoms are nodes, and their connections represent participation in the same topological feature. This network then provides quantifiable characteristics that serve as input features for our machine learning models, effectively translating complex topological information into biologically relevant descriptors.

**Molecular Docking** ATP^4-^ and peptide substrates were docked sequentially to the relaxed ALK, BRAF, EGFR, HER2, and MEK1 active KD structures extracted from the last frame of the MD simulations with two Mg^2+^ ions using Schrodinger’s (v. 2020.3) Induced-Fit Docking (IFD) [18] and Glide protocols [19] Positions of the Mg^2+^ ions were modeled from Mn^2+^ ions in a ternary PKACA-ATP-Peptide complex (PDB ID: 1ATP) after aligning the receptor KDs to the reference structure. In addition, water molecules within 10Å of the Mg^2+^ ions were retained throughout the docking simulations.[20] The proposed peptide substrates for RTKs were modeled from the autophosphorylation sites on the RTKs themselves, namely Y1278 from the Y’XXX’YY autophosphorylation motif on the A-loop of ALK TKD and the prominent autophosphorylation site Y1092 (PEYINQ) of the cytoplasmic tail region of EGFR [21, 22] As HER2 heterodimerizes with EGFR for its activation, the same peptide substrate for EGFR TKD was used. [23] For the Ser/Thr-protein kinases BRAF and MEK1, phosphorylation sites from the A-loop of downstream effectors were used as peptide substrates. Specifically, the peptide substrates were modeled from S218 (IDSMA, PDB ID: 3EQD) of MEK1 and T183/Y185 (FLTEYVA, PDB ID: 5UMO) of ERK2 [24, 25]

Ten stereoisomers were generated and clustered for ATP^4-^ before being docked to the receptor using the IFD protocol with the OPLS3e forcefield. [26] A 20Å box was defined using the centroid of the Asp, Lys, and Gly residues from the DFG motif, KE salt bridge, and the P-loop, respectively. [27] All other settings in the ATP IFD simulations were kept default. From the resulting poses of the KD-ATP complex, poses with the lowest docking score from each cluster were selected to be the receptor structure of peptide docking. [28] With the γ-phosphate in the center of a 25Å simulation box, receptor grids were generated for peptide docking, and rotation was allowed for hydroxyl and thiol groups inside the box. A constraint was implemented to ensure that at least one atom from the peptide substrates was within 4Å of the γ-phosphate donor to sample KD-ATP-peptide poses in which the transition state of nucleophilic attack is present. The proposed peptide substrates were subsequently docked to the receptor grids using the Glide protocol. Poses were scored and ranked using the GlideScore function, which approximates the binding affinity. The top-ranked poses were filtered by assessing their biological validity, where the -OH group on the side chain of the substrate phosphorylation sites was oriented toward the ATP γ-phosphate [4]. From the valid pose with the lowest peptide docking score, the binding pocket of the receptor was determined by selecting full residues within 6Å of the centroid of the bound ATP^4-^ and peptide substrate, as well as residues from the DFG motif, the HRD motif, and the KE salt-bridge [11]. Using heavy atoms in the binding pocket-ATP^4-^-Mg^2+^-peptide complexes, the topology of the ternary complex was constructed.

**Generation of docking feature.** Wet docking was performed to generate ternary complexes for 215 ALK, BRAF, EGFR, HER2, and MEK1 variants. Water molecules and one Mg^2+^, labeled MG1, were modeled from a ternary complex of the active IRK structure (PDB ID: 3BU5). The position of a second Mg^2+^ (MG2) was modeled from that of the Mn^2+^ in a receptor-ATP-peptide system of PRKACA (PDB ID: 1ATP). Using the top-ranked valid ternary complexes of the variant systems, the interatomic distances between key atoms in the phosphorylation transition state were measured and used as features in training. These distances included:

1. Distance between 𝛾-phosphate and the peptide phosphorylation site side chain O atom
2. Distance between MG1 and MG2
3. Distance between 𝛽-phosphate and MG1
4. Distance between 𝛾-phosphate and MG1
5. Distance between DFG-Asp C-gamma and MG2
6. Distance between HRD-Asp C-gamma and MG2
7. Distance between HRD-Asp C-gamma and the peptide phosphorylation site side chain O atom

In addition, binding pockets of the KDs were determined by accounting all residues within a sphere centered at the geometric center of the ATP and peptide substrate in their active wild-type systems. Two radii of the sphere were defined for the binding pockets: the default radius used in Schrodinger Maestro [18,19] for the determination of binding pockets, and the smallest radius where the DFG and HRD motifs are fully included in the binding pockets. The determined large and small binding pockets were used to generate topology constructs and feature generation.

**Baseline machine learning methods**

Support vector machine (SVM), gradient boosting decision tree (GBDT), and natural gradient boosting decision tree (nGBDT) are used to predict the activation status for kinases. Docking and MD features are considered as privileged features. The rest features are considered as basic features. As not all kinases have docking and MD features, we use K nearest neighbor imputation (implemented by scikit-learn) to impute the missing docking and MD features. SVM, GBDT and K nearest neighbor imputation are implemented by scikit-learn (K is set to $\sqrt{n}$, where $n$ is the size of the kinase dataset).

**Memory-based neural network**

Let $d_{basic}$, and $d_{privileged}$ denote the dimensions of basic and privileged features, respectively. We define $D_{p}=\{x_{i, basic}^{p},x_{i,privileged}^{p},y_{i}^{p}|i=1,2,...,n_{p}, x_{i, basic}^{p} \in R^{d_{basic}}, x_{i, privileged}^{p} \in R^{d_{privileged}}\}$ to be the kinase dataset whose samples have the privileged information (i.e., docking and MD features), and $y_{i}^{p}$ stands for the activation label of sample $x_{i}^{p}$. Similarly, we define $D_{c}=\{x_{i, basic}^{c},y_{i}^{c}| i=1,2,...,n_{c}, x_{i}^{c} \in R^{d_{basic}}\}$ to be the kinase dataset whose samples do not have the privileged information.

We first describe how our neural network method predicts the activation labels for $D_{p}$ and then show how it can be extended to do the same task for $D_{c}$. Let $MLP()$ be a fully connected neural network, our neural network predictor uses a $MLP_{main}()$ with $L$ layers of nonlinear transformations to process the basic features of $x_{i, basic}^{p}$

${h^{p}}_{i, basic,1}, {h_{i,basic,2}}^{p}, ...,{h_{i,basic,L}}^{p} =MLP_{main}(x_{i, basic}^{p}$),

where ${h^{p}}_{i, basic,j}$ ($j$ from 1 to $L$) stands for the output of $x_{i,basic}^{p}$ from the layer of $j$ of $MLP_{main}()$. ${h^{p}}_{i, basic,j}$ will be further adjusted by the features extracted from the privileged information before it is be inputted into the $\left( j+1 \right)th$ layer of $MLP_{main}()$.

We then define the process of extracting features from the privileged information. To this end, we use a $MLP_{p1}()$ to first process the concatenation of privileged and basic features for $x_{i}$

$f_{i}^{p} = MLP_{p1}(concat\left( x_{i, basic}^{p}, x_{i, privileged}^{p} \right))$.

Sinusoidal positional encoding is then used to create two vectors $POS_{shift, j}$ and $POS_{scale, j}$ with the same size as $f_{i}^{p}$ , and $j$ denotes the layer index of $MLP_{main}()$. We used two single-layer MLPs ($MLP_{aux1}()$ and $MLP_{aux2}()$) to transform $POS_{shift, j}$ and $POS_{scale, j}$, and then perform the element-wise shift and scaling operation on $f_{i}^{p}$ to create layer number dependent ${f^{p}}_{i,j}$:

${f^{p}}_{i,j} = MLP_{aux1}(POS_{shift, j}) + MLP_{aux2}(POS_{scale, j}) *f_{i}^{p}$.

We then use another two MLPs ($MLP_{p2}()$ and $MLP_{p3}()$) to further process ${f^{p}}_{i,j}$and generate the shift and scaling factors to adjust ${h^{p}}_{i,basic,j}$:

${h^{p,adjusted}}_{i,basic,j}$ = $MLP_{p2}({f^{p}}_{i,j}) + MLP_{p3}({f^{p}}_{i,j}) *{h^{p}}_{i, basic,j}$.

Here, ${h^{p,adjusted}}_{i,basic,j}$ is the adjusted hidden features of $x_{i}^{p}$ extracted from the $j$th layer of $MLP_{main}()$ and adjusted by privileged information of $x_{i}^{p}$. It is also the input of $MLP_{main}()$ in $(j+1$)th layer.

To enable the $x_{z}^{c}\in D_{c}$ to be adjusted by the privileged information, we adopted a memory-based strategy. Specifically, we store all ${h^{p}}_{i, basic,j}$, $MLP_{p2}({f^{p}}_{i,j})$ and $MLP_{p3}({f^{p}}_{i,j})$for $x_{i}^{p}\in D_{p}$. For $x_{z}^{c}$, we again use $MLP_{main}()$ to produce the intermediate features

${h^{c}}_{z, basic,1}, {h_{z,basic,2}}^{c}, ...,{h_{z,basic,L}}^{c} =MLP_{main}(x_{z, basic}^{c}$).

The scaling and shift factors of ${h^{c}}_{z, basic,j}$ are generated by the following attention mechanism [29]

${SHIFT}_{z,j}= \sum_{=1,...,n_{p}} \alpha_{i,j}^{z}*MLP_{p2}({f^{p}}_{i,j})$, $SCALING_{z,j}=\sum_{=1,...,n_{p}} \alpha_{i,j}^{z}*MLP_{p3}({f^{p}}_{i,j})$,

where $\alpha_{i,j}^{z}$= $\frac{e^{<{h^{c}}_{z, basic,j}，{h^{p}}_{i, basic,j}>}}{\sum_{=1,...,n_{p}} e^{<{h^{c}}_{z, basic,j}，{h^{p}}_{i, basic,j}>}}$ (<> stands for the inner product).

With this imputed scaling and shift factors, we can have,

${h^{c, adjusted}}_{z,basic,j} = {SHIFT}_{z,j}+ SCALING_{z,j}*{h^{c}}_{z, basic,j}$ .

For $MLP_{main}$, $MLP_{p1}$, $MLP_{p2}$ and $MLP_{p3}$, a single layer sequentially consists of a linear transformation, a layer normalization, a dropout layer and a Gaussian Error Linear Unit (GELU) layer. We also constrain $MLP_{p2}$ and $MLP_{p3}$ to share the same weights expect their last layers. For $MLP_{aux1}$ and $MLP_{aux2}$, they have one single layer and is constructed similarly except not having the layer normalization.

After obtaining ${h^{p, adjusted}}_{i,basic,L}$ or ${h^{c, adjusted}}_{i,basic,L}$ for $x_{i}$ from $D_{p}$ or $D_{c}$, respectively, we consider it as a feature representation of $x_{i}$ and use Deep Neural Decision Forests [30] to make predictions for kinase activation. Specifically, for a neural decision tree with a depth of $k$, it is a complete binary tree with $2^{k}$ and $2^{k}-1$ leaf nodes and internal nodes, respectively. This neural decision tree randomly selects $r\%$ features from ${h^{p, adjusted}}_{i,basic,L}$ (or ${h^{c, adjusted}}_{i,basic,L}$) to use a linear transformation followed by a sigmoid function to map the selected features to $2^{k}-1$ values, which are interpreted as the soft routing probabilities (i.e., go to left or right child node) of the data instance $x_{i}$ on the $2^{k}-1$ internal nodes. Using these routing probabilities, we can determine the probability of $x_{i}$ reaching each leaf node from the root of the decision tree. Each leaf node is considered as a decision-making node for binary classification. The class logits are randomly initialized and will be learnt from the data. The final prediction of the neural decision forest is computed as the weighted sum of the softmax-normalized logits of each leaf node, where the weights correspond to the probability of reaching that leaf across all nodes in all decision trees. We trained our memory-based neural network with focal loss. [31] To enable a robust prediction, we also train five different model copies using different random seeds and average the prediction results.

**Computational evaluation and hyperparameter tuning**

We conducted five repeated five-fold cross validation (CV) under different random seeds and reported the average prediction performances (e.g., balanced accuracy [BACC], area under precision-recall curve [AUPR], area under receiver operating characteristic curve [AUROC]) and corresponding standard deviation. For each machine learning model, we used grid search to search hyperparameter combinations and select the best performing model based on AUROC under our CV scheme. Classic machine learning models were implemented by scikit-learn while neural networks were implemented by pytorch. Here, we reported our hyperparameter search ranges:

Tree-based models: we select the number of trees from [32, 128, 512], and the maximum tree depth from [4, 8, 16, 32, None].

For the SVM: we select C from [10, 1, 0.1, 0.01].

Neural network: we select dropout in neural network from [0.7, 0.6, 0.5, 0.4, 0.3, 0.2, 0.1], ratio r from the neural decision forest from [0.8, 0.7, 0.6, 0.5, 0.4, 0.3, 0.2, 0.1], depth of the neural decision tree from [9, 8, 7, 6, 5, 4, 3, 2], number of neural decision tree from [512, 256, 128], gamma in the focal loss from [12, 10, 8, 5, 3], training epoch from [3000, 2500, 2000, 1500, 1000, 900, 800, 700, 600, 500, 400, 300], $MLP_{main}$’s hidden layer sizes from [[512], [512, 512], [512, 512, 512], [256], [256, 256], [256, 256, 256], [128], [128, 128], [128, 128, 128]] (where the length of a list is the number of hidden layers, and the numbers in the list are corresponding numbers of hidden units), $MLP_{p1}$’s, $MLP_{p2}$’s and $MLP_{p3}$’s hidden layer sizes from [[512], [512, 512], [256], [256, 256], [128], [128, 128]]. We used full batch gradient descent to train the model. We selected AdamW [32] as the optimizer with the following hyperparameters lr=0.001, betas=(0.9, 0.98), eps=1e-8, weight_decay=0.05. We also used a learning rate scheduler get_linear_schedule_with_warmup in pytorch with num_warmup_steps=50. We clipped the gradient norm when it is over 1.

Feature importance**:** for the GDBT models, we used permutation feature importance to derive the importance scores for features. For neural networks, integrated gradient was used to derive the feature importance. Specifically, given a kinase mutation, the baseline of it for the integrated gradient approach was defined as the features of its corresponding wild type.

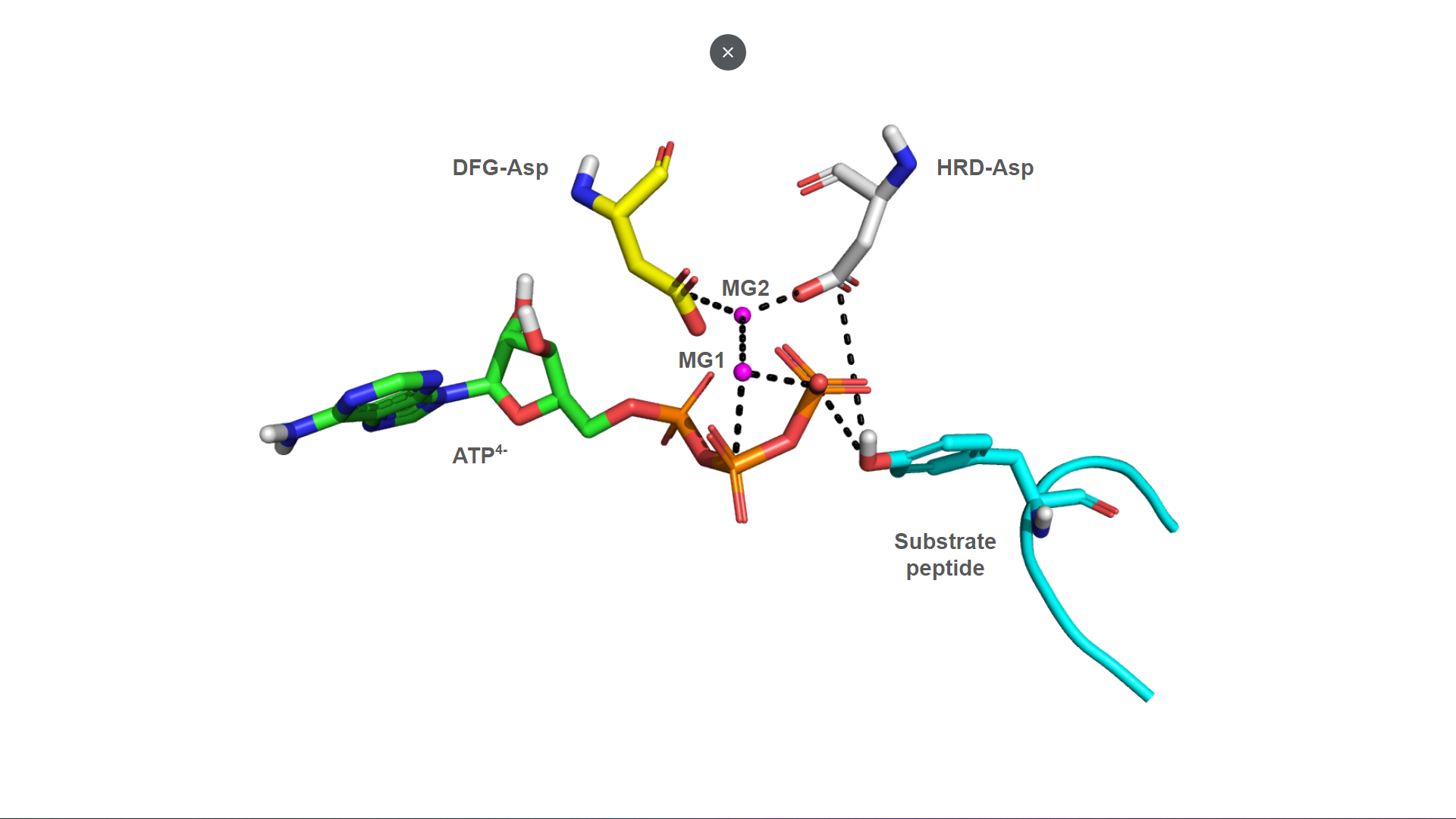

**Fig. S1.** The ternary complex of KD-ATP-peptide (only the Asp in conserved motifs HRD and DFG were shown for the KD. Black dashed lines denote the measured interatomic distances between atom pairs.

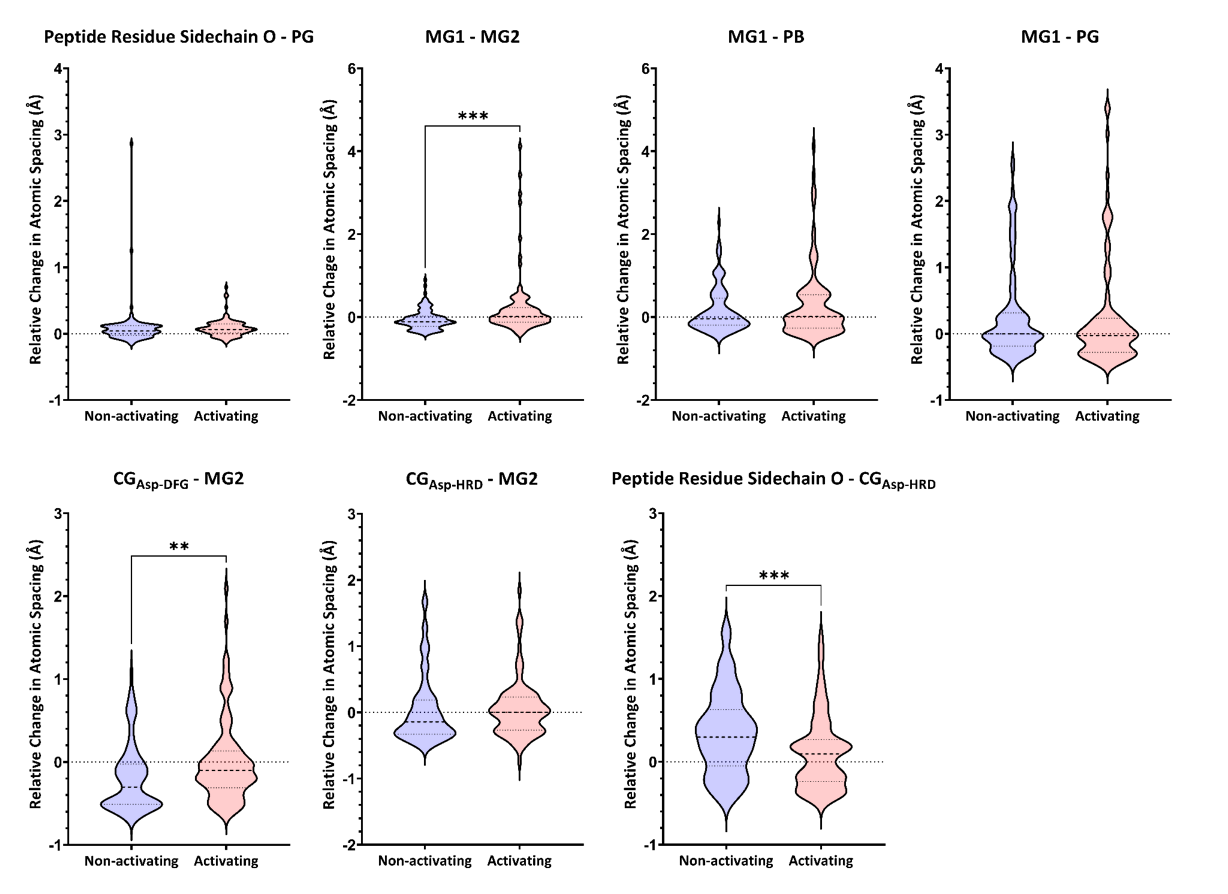

**Fig. S2.** **Violin plots (and statistical analysis) of the distance-based features.** Distance #2,5,7 resulted in statistically significant differences between activating and non-activating mutations in terms of T-test (mean); Distances #1,2,3,5 resulted in statistically significant differences in terms of F-test (distribution). Pearson correlation coefficients were calculated between the distances and the labels. Coefficients of distances #2 (~-0.35) and #5 (~0.25). Note: The distance features derived from the peptide-receptor poses, using the **relative changes in the distances (using distances in WT poses as reference)** as inputs yielded **better NN performance in terms of BACC**. It was observed that when the relative changes were paired with feature groups from the basic feature pool, the BACC increased by a considerable amount (>3% when paired with the whole feature pool, >6% when paired with different biochemical feature groups). These features were also tested along with the privileged feature groups (MD, protein backbone topology) and also resulted in a slight increase in the NN BACC. In addition, these 7 new features alone could give a ~63% NN BACC with 215 samples.

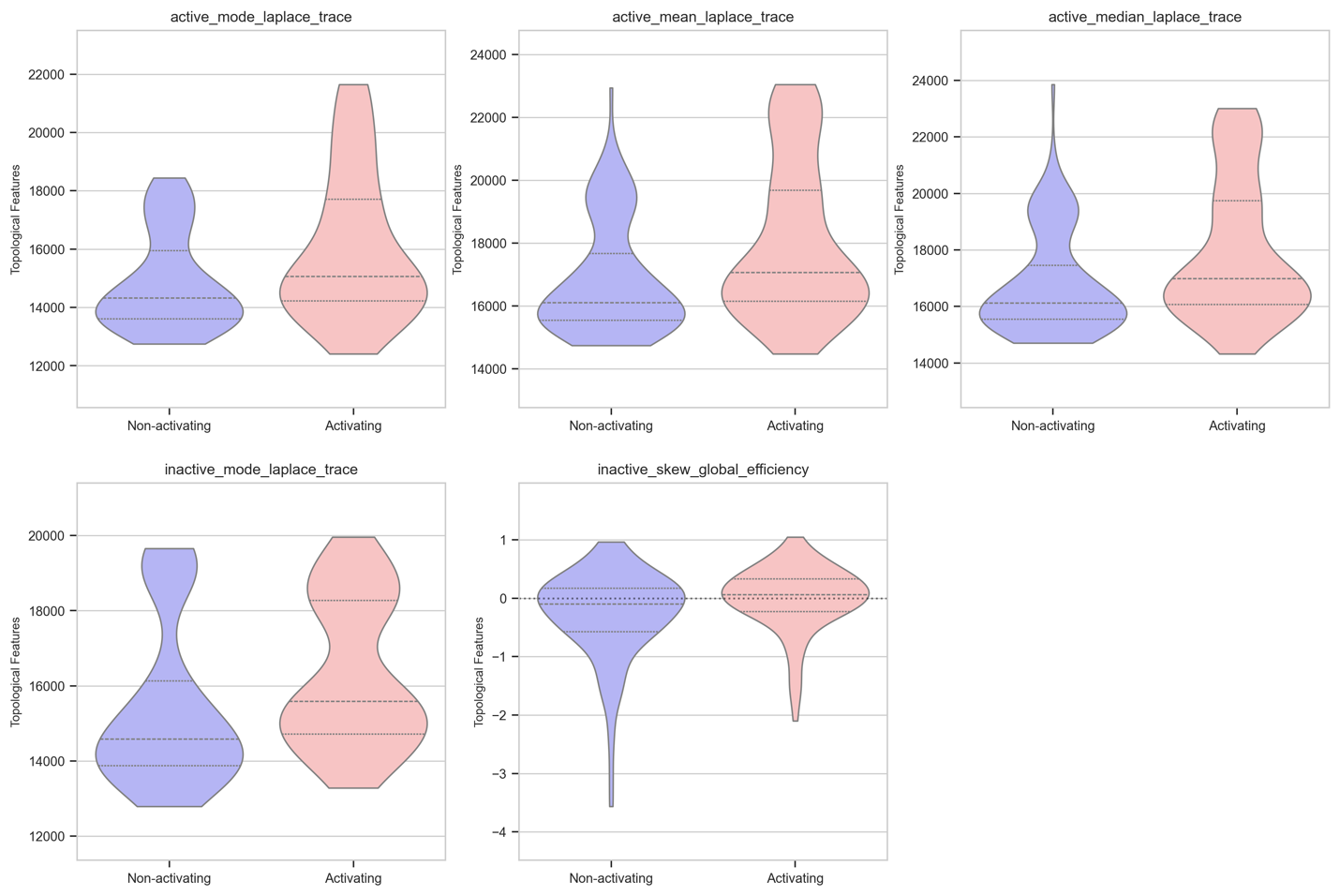
**Fig. S3.** Violin plots of top 5 topological features that are selected based on the Pearson correlation coefficient with the activation label of 215 kinase mutants.

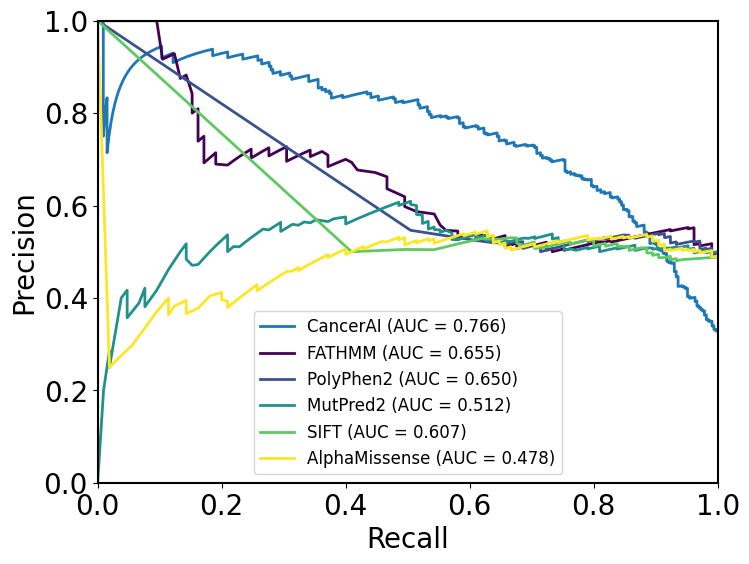

**Fig. S4.** Comparison of recall precision characteristic (PRC) curves of Kinome-AI (averaged values from 5-fold cross validation), FATHMM, Polyphen2, MutPred2, SIFT, and AlphaMissense using 21% of total 1003 kinase mutants. The area under the curve (AUC) indicated in parenthesis. A higher AUC score indicates better performance.

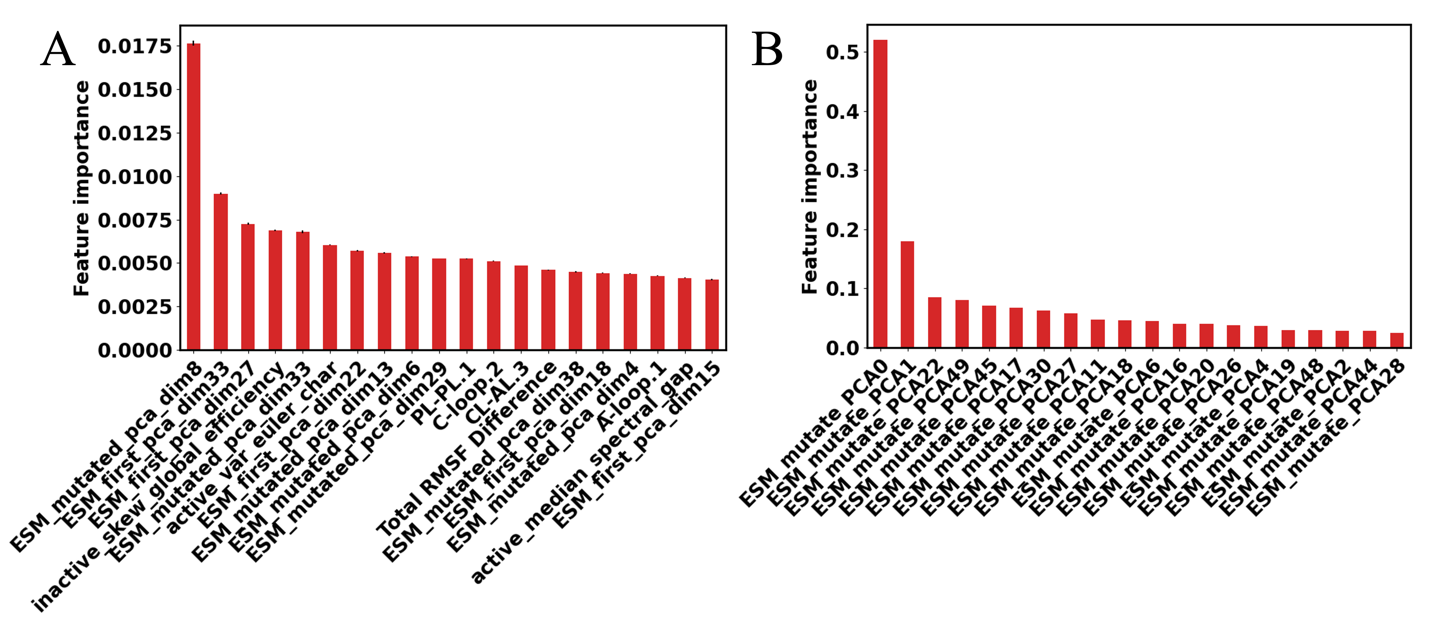

**Fig. S5.** Top 20 selected structural and topological features according to the permutation feature importance of the (A) GBDT model and from (B) integrated gradient method of NN-based model.

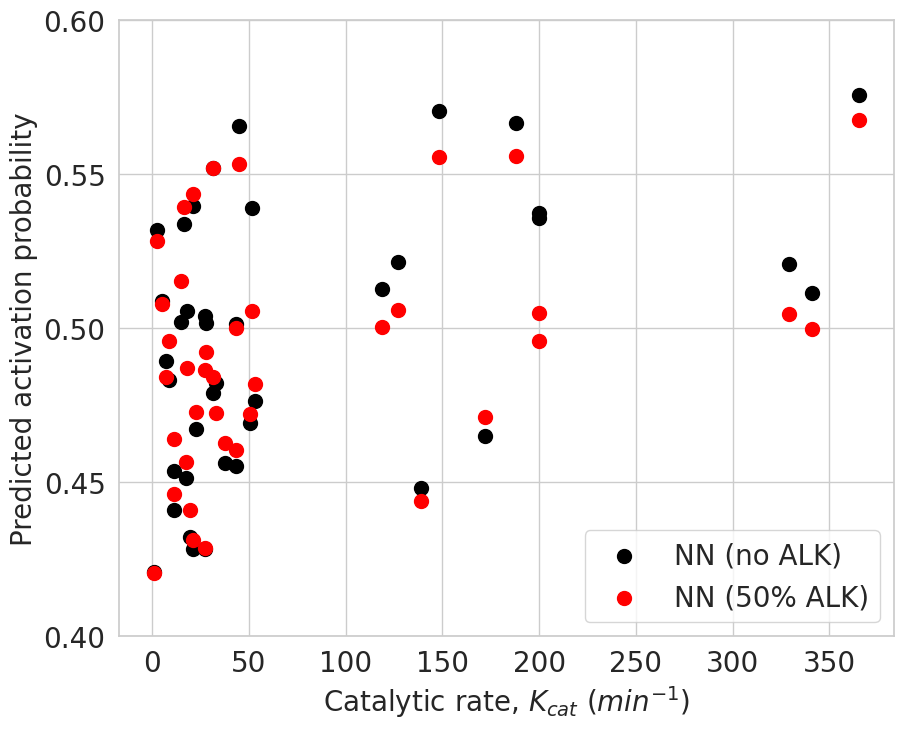

**Fig. S6.** Correlation between Kinome-AI predicted activation probability of ALK mutants and experimental catalytic rate (k_cat_) taken from Patil et al [27]. Black and red dots represent the prediction based on retraining model without all 42 ALK mutants and with random selected 50% of 42 ALK mutants, respectively, in which case the corresponding Pearson correlation coefficients are 0.343 and 0.437.

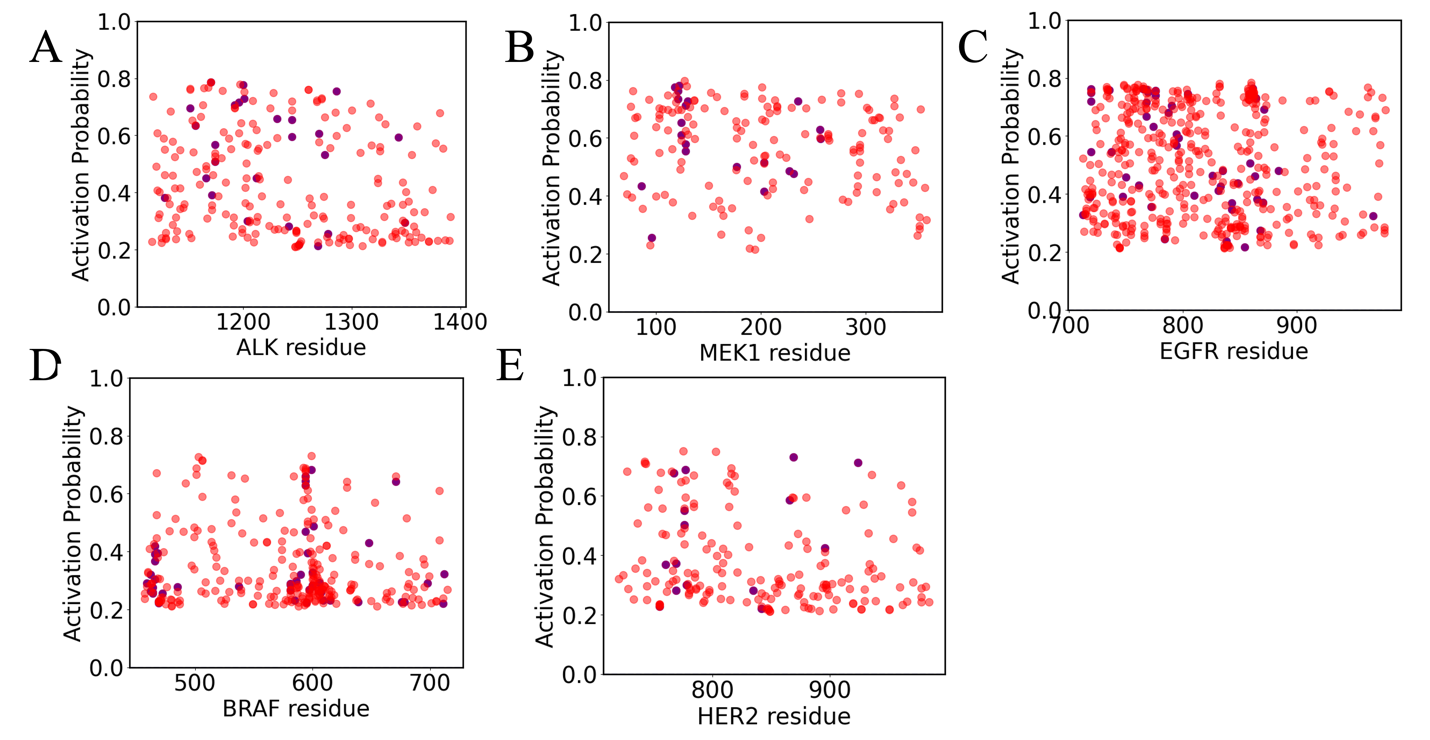

**Fig. S7.** Prediction of activation probability of (A) ALK, (B) MEK1, (C) EGFR, (D) BRAF and (E) HER2 kinase mutations taken from the COSMIC dataset (<https://cancer.sanger.ac.uk/cosmic>).

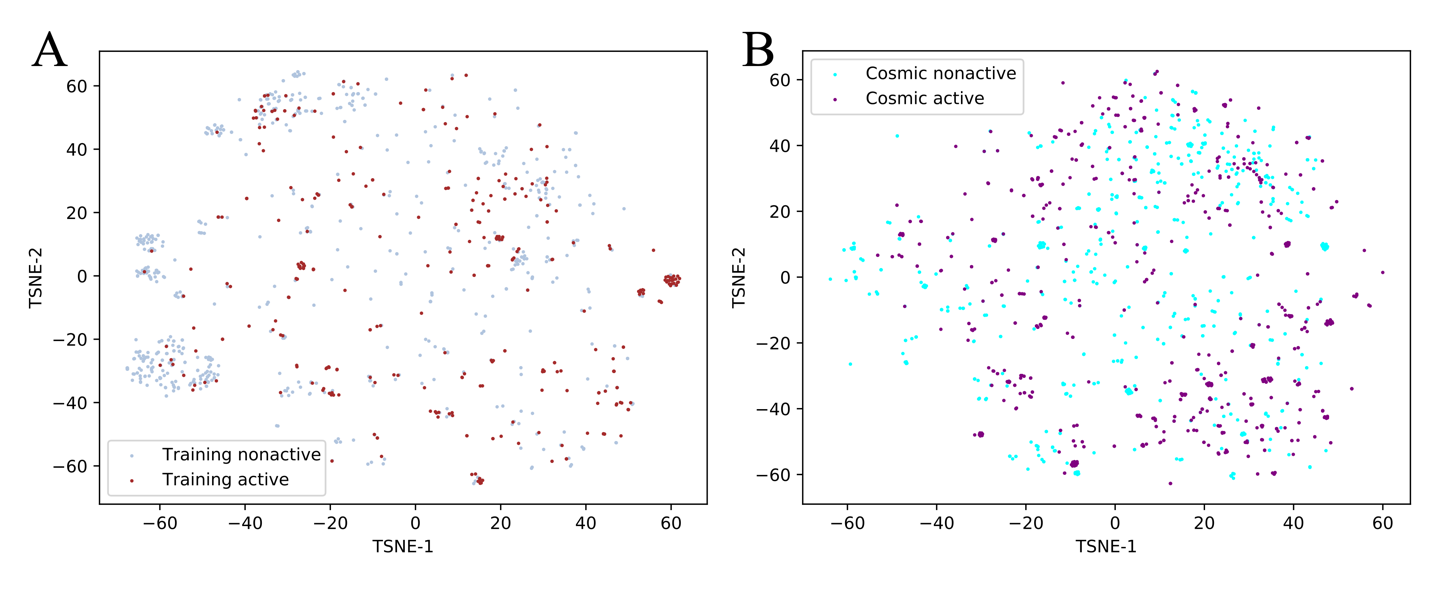
**Fig. S8.** Clustering visualization plot of kinase mutants (A) our dataset and (B) COSMIC database using both the first and mutation site token of kinase domain ESM2 features using the nonlinear dimensionality reduction methods (T-SNE) [33]. The color scheme assigned according to the predicted active and non-active labels using neural network model).

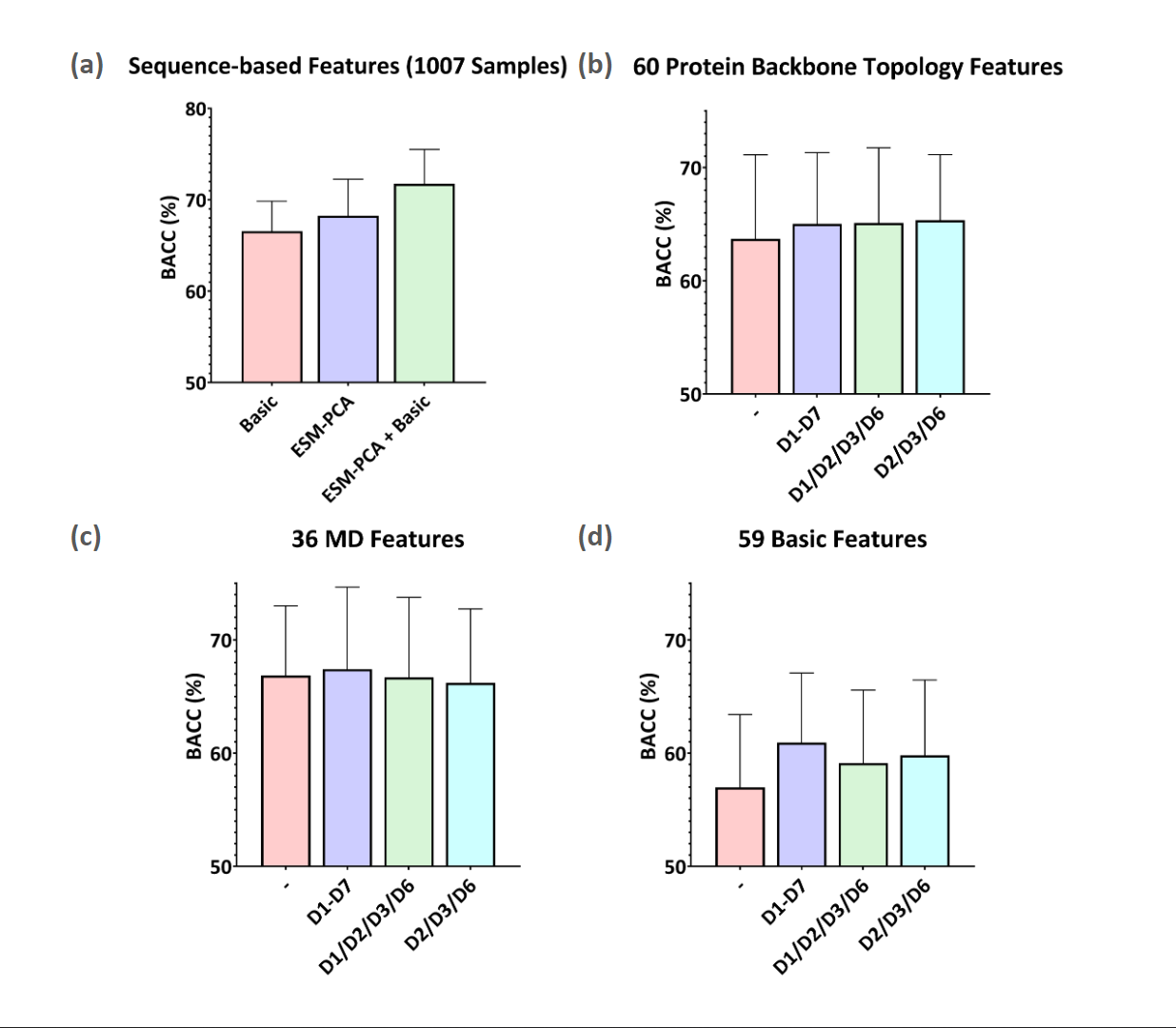

**Fig. S9.** (a) Comparison of ML performance using biochemical and ESM features for all 1003 mutants (b)-(d) Comparison of ML performance using biochemical, ESM features, and MD features for 221 mutants. Each feature group was paired with different distance-based docking features to evaluate the performance of the docking features.

**Table S1.** Selected peptide substrates for kinase docking. Ser/Thr/Tyr on the peptides are either autophosphorylation sites identified on the kinases or phosphorylation sites on their downstream effectors [34]. The EGFR and HER2 kinase domains share the same peptide substrate due to the heterodimer formation for signal transduction [35].

Note: ^**^For EGFR/HER2, the proposed peptide PEY^1092^IN is on the C-terminal tail of EGFR. As crystal structures of the tail are not available, the peptide was modeled and mutated from a HER2 peptide ^1137^PEY**V**NQ^1142^ in PDB ID:2L4K.

*PDB ID:2GS6 presents the active EGFR in complex with a Src substrate peptide EIYGE.

| Kinase Domains | Selected Peptide Substrate | Modeled from |
| --- | --- | --- |
| ALK | ^*^DIY^1278^RA | Src substrate peptide EIYGE in PDB ID:2GS6 |
| BRAF | IDS^218^MA | Phosphorylation site of downstream effector MEK1, PDB ID: 3EQD |
| EGFR  HER2 | ^**^PEY^1092^IN | Autophosphorylation site Tyr1092 on EGFR, PDB ID: 2L4K |
| MEK1 | FLT^183^EY^185^VA | Phosphorylation sites of downstream effector ERK2, PDB ID: 5UMO |

**Table S2.** The table below summarizes all features used in the machine learning algorithm training.

| Feature Category/Group | | Features | Counts |
| --- | --- | --- | --- |
| Sequence-based | Bioinformatics and Biochemical | Wild-type residue^*^ | 20 |
|  |  | Wild-type residue type^*^ | 5 |
|  |  | Mutated residue^*^ | 20 |
|  |  | Mutated residue type^*^ | 5 |
|  |  | Location of mutation^*^ | 4 |
|  |  | Normalized difference between mutation and wild-type biochemical properties | 5 |
|  | ESM Embeddings | PCA-reduced token of wild-type | 100 |
|  |  | PCA-reduced token of mutant | 100 |
| Structure-based | Molecular Dynamics | ΔRMSF | 12 |
|  |  | ΔSASA | 12 |
|  |  | ΔRGYR | 10 |
|  |  | A_MUT_ | 2 |
|  |  | Pairwise subdomain interactions | 40 |
|  | Molecular Docking | Relative change in atom-pair distances in the phosphorylation transition state | 7 |
|  | Topology | Protein backbones | 60 |
|  |  | Active site heavy atoms | 60 |
|  |  | Docking-derived binding pocket heavy atoms | 5 |

^*^ Binary features.

**Table S3**. Comparison of binary classification performance of different ML methods using all features except the ESM-2 features. 5-fold cross validation (repeated five times, mean values were reported).

| **Models** | **AUROC** | **AUPR** | **BACC** |
| --- | --- | --- | --- |
| **Memory-based NN+Tree** | **0.797 ± 0.007** | 0.620 ± 0.011 | **0.725 ± 0.004** |
| GBDT (knn imputation) | 0.784 ± 0.008 | 0.633 ± 0.012 | 0.682 ± 0.009 |
| **GBDT (no imputation)** | 0.786 ± 0.005 | **0.635 ± 0.006** | 0.693 ± 0.008 |
| nGBoost (knn imputation) | 0.781 ± 0.007 | 0.614 ± 0.014 | 0.657 ± 0.012 |
| nGBoost (no imputation) | 0.785 ± 0.010 | 0.628 ± 0.013 | 0.652 ± 0.013 |
| SVM (knn imputation) | 0.783 ± 0.008 | 0.595 ± 0.007 | 0.666 ± 0.008 |
| SVM (no imputation) | 0.778 ± 0.008 | 0.588 ± 0.009 | 0.646 ± 0.009 |

**Table S4**. Comparison of binary classification performance of different ML methods using all features except the ESM-2 features. 5-fold cross validation (repeated five times, mean values were reported).

| Data with docking and MD features | **Models** | **AUROC** | **AUPR** | **BACC** |
| --- | --- | --- | --- | --- |
|  | **Memory-based NN+Tree** | 0.703 ± 0.013 | 0.647 ± 0.013 | **0.678 ± 0.014** |
|  | GBDT (knn imputation) | 0.728 ± 0.025 | 0.708 ± 0.022 | 0.660 ± 0.022 |
|  | **GBDT (no imputation)** | **0.734 ± 0.013** | **0.718 ± 0.017** | 0.666 ± 0.019 |
|  | nGBoost (knn imputation) | 0.709 ± 0.027 | 0.682 ± 0.039 | 0.632 ± 0.023 |
|  | nGBoost (no imputation) | 0.705 ± 0.035 | 0.677 ± 0.032 | 0.627 ± 0.038 |
|  | SVM (knn imputation) | 0.710 ± 0.018 | 0.680 ± 0.031 | 0.613 ± 0.014 |
|  | SVM (no imputation) | 0.717 ± 0.025 | 0.659 ± 0.045 | 0.580 ± 0.019 |
| Data without docking and MD features | **Models** | **AUROC** | **AUPR** | **BACC** |
|  | **Memory-based NN+Tree** | **0.807 ± 0.007** | **0.610 ± 0.011** | **0.723 ± 0.007** |
|  | GBDT (knn imputation) | 0.784 ± 0.006 | 0.595 ± 0.013 | 0.673 ± 0.007 |
|  | GBDT (no imputation) | 0.786 ± 0.005 | 0.604 ± 0.009 | 0.686 ± 0.009 |
|  | nGBoost (knn imputation) | 0.783 ± 0.007 | 0.579 ± 0.008 | 0.645 ± 0.011 |
|  | nGBoost (no imputation) | 0.791 ± 0.005 | 0.603 ± 0.011 | 0.644 ± 0.014 |
|  | SVM (knn imputation) | 0.790 ± 0.010 | 0.575 ± 0.010 | 0.691 ± 0.007 |
|  | SVM (no imputation) | 0.783 ± 0.009 | 0.567 ± 0.009 | 0.674 ± 0.009 |

**Table S5**. Comparison of binary classification performance of different ML methods using all features. 5-fold cross validation (repeated five times, mean values were reported).

| **Models** | **AUROC** | **AUPR** | **BACC** |
| --- | --- | --- | --- |
| **Ensemble** | **0.849 ± 0.003** | **0.747 ± 0.005** | **0.763 ± 0.003** |
| **Memory-based NN+Tree** | 0.842 ± 0.004 | 0.732 ± 0.009 | **0.763 ± 0.005** |
| **GBDT (knn imputation)** | **0.843 ± 0.007** | 0.736 ± 0.008 | 0.755 ± 0.006 |
| **GBDT (no imputation)** | 0.840 ± 0.006 | **0.738 ± 0.013** | 0.753 ± 0.005 |
| nGBoost (knn imputation) | 0.816 ± 0.006 | 0.677 ± 0.010 | 0.705 ± 0.007 |
| nGBoost (no imputation) | 0.812 ± 0.011 | 0.674 ± 0.019 | 0.700 ± 0.014 |
| SVM (knn imputation) | 0.825 ± 0.008 | 0.702 ± 0.007 | 0.717 ± 0.006 |
| SVM (no imputation) | 0.839 ± 0.005 | 0.736 ± 0.012 | 0.751 ± 0.007 |

Note: (1). Ensemble means averaged results combining the NN and GBDT (knn imputation) model (2). The memory-based NN model doesn’t use basic features as the performance decreased by including basic features.

**Table S6**. Comparison of binary classification performance of different ML methods using all features. 5-fold cross validation (repeated five times, mean values were reported).

| Data with docking and MD features | **Models** | **AUROC** | **AUPR** | **BACC** |
| --- | --- | --- | --- | --- |
|  | **Ensemble** | **0.791 ± 0.017** | **0.759 ± 0.023** | **0.726 ± 0.025** |
|  | **Memory-based NN+Tree** | 0.773 ± 0.008 | **0.757 ± 0.015** | **0.719 ± 0.020** |
|  | **GBDT (knn imputation)** | **0.783 ± 0.025** | 0.750 ± 0.024 | 0.716 ± 0.023 |
|  | GBDT (no imputation) | 0.773 ± 0.019 | 0.765 ± 0.024 | 0.709 ± 0.027 |
|  | nGBoost (knn imputation) | 0.749 ± 0.010 | 0.723 ± 0.011 | 0.682 ± 0.012 |
|  | nGBoost (no imputation) | 0.727 ± 0.014 | 0.704 ± 0.026 | 0.656 ± 0.014 |
|  | SVM (knn imputation) | 0.724 ± 0.023 | 0.669 ± 0.036 | 0.624 ± 0.013 |
|  | SVM (no imputation) | 0.776 ± 0.012 | 0.752 ± 0.024 | 0.710 ± 0.013 |
| Data without docking and MD features | **Models** | **AUROC** | **AUPR** | **BACC** |
|  | **Ensemble** | **0.85 ± 0.003** | **0.734 ± 0.007** | **0.758 ± 0.005** |
|  | **Memory-based NN+Tree** | **0.847 ± 0.004** | 0.718 ± 0.007 | **0.764 ± 0.003** |
|  | GBDT (knn imputation) | 0.843 ± 0.006 | 0.723 ± 0.009 | 0.751 ± 0.006 |
|  | GBDT (no imputation) | 0.844 ± 0.008 | 0.722 ± 0.016 | 0.754 ± 0.007 |
|  | nGBoost (knn imputation) | 0.818 ± 0.008 | 0.652 ± 0.018 | 0.691 ± 0.007 |
|  | nGBoost (no imputation) | 0.818 ± 0.013 | 0.657 ± 0.026 | 0.693 ± 0.015 |
|  | SVM (knn imputation) | 0.835 ± 0.010 | 0.708 ± 0.009 | 0.753 ± 0.005 |
|  | **SVM (no imputation)** | 0.842 ± 0.004 | **0.726 ± 0.009** | 0.750 ± 0.007 |

Note: 1. The memory-based NN+Tree model doesn’t use basic features as the performance decreased by including basic features; 2. Ensemble stands for the averaged results of NN and GBDT (knn imputation) models.

**Table S7**. Comparison of binary classification performance of two best ML methods on the ALK mutants reported in Table 2 of Patil et al [27].

| **GBDT model** | **Binarization threshold** | **True positive rate (TPR)** | **False positive rate (TPR)** | **Balanced accuracy (BACC)** |
| --- | --- | --- | --- | --- |
|  | 0.5 | 0.944 | 0.750 | 0.597 |
|  | 0.6 | 0.889 | 0.708 | 0.590 |
|  | 0.7 | 0.833 | 0.625 | 0.604 |
|  | 0.8 | 0.722 | 0.500 | 0.611 |
|  | 0.9 | 0.667 | 0.208 | 0.729 |
| **Memory-based NN model** | **Binarization threshold** | **True positive rate (TPR)** | **False positive rate (TPR)** | **Balanced accuracy (BACC)** |
|  | 0.2 | 1.0 | 1.0 | 0.5 |
|  | 0.3 | 1.0 | 1.0 | 0.5 |
|  | 0.4 | 1.0 | 1.0 | 0.5 |
|  | 0.5 | 0.611 | 0.25 | 0.681 |
|  | 0.6 | 0.0 | 0.0 | 0.5 |

Note: Model are retrained without ALK mutants.

**Table S8. Prediction performance comparison of GBDT, MNN and the ensemble of GBDT and MNN models on 8 different kinase classes. Kinase classification notations refer to Fig. 1A. Results are averaged over five different random seeds.**

| **Kinase class** | **# mutants (N_total_=1003)** | **Balanced accuracy** | | |
| --- | --- | --- | --- | --- |
|  |  | **GBDT** | **MNN** | **GBDT MNN ensemble** |
| **TK** | **427** | 0.730 ± 0.011 | 0.753 ± 0.004 | 0.740 ± 0.012 |
| **TKL** | **170** | 0.636 ± 0.015 | 0.639 ± 0.023 | 0.645 ± 0.008 |
| **STE** | **90** | 0.884 ± 0.020 | 0.806 ± 0.016 | 0.881 ± 0.020 |
| **CK1** | **4** | 1.000 ± 0.000 | 1.000 ± 0.000 | 1.000 ± 0.000 |
| **AGC** | **51** | 0.717 ± 0.059 | 0.817 ± 0.044 | 0.706 ± 0.054 |
| **CAMK** | **99** | 0.648 ± 0.048 | 0.587 ± 0.003 | 0.650 ± 0.047 |
| **CMGC** | **77** | 0.593 ± 0.030 | 0.496 ± 0.056 | 0.576 ± 0.006 |
| **Other** | **85** | 0.573 ± 0.038 | 0.579 ± 0.042 | 0.573 ± 0.038 |

**Table S9**. Definition of starting and ending residue of kinase sub-domains.

| Kinase name | Kinase domain | | Phosphate binding loop | | αC helix | | Catalytic loop | | Activation loop | |
| --- | --- | --- | --- | --- | --- | --- | --- | --- | --- | --- |
|  | start | end | start | end | start | end | start | end | start | end |
| ALK | 1116 | 1392 | 1123 | 1128 | 1158 | 1176 | 1243 | 1254 | 1268 | 1291 |
| BRAF | 457 | 717 | 464 | 469 | 491 | 509 | 570 | 581 | 592 | 615 |
| EGFR | 712 | 979 | 719 | 724 | 752 | 770 | 831 | 842 | 853 | 876 |
| MEK1 / MAP2K1 | 68 | 381 | 75 | 81 | 107 | 125 | 185 | 194 | 208 | 233 |
| ERBB2 / HER2 | 720 | 987 | 727 | 732 | 760 | 778 | 839 | 850 | 861 | 884 |
